# Towards a unified theory of plant photosynthesis and hydraulics

**DOI:** 10.1101/2020.12.17.423132

**Authors:** Jaideep Joshi, Benjamin D. Stocker, Florian Hofhansl, Shuangxi Zhou, Ulf Dieckmann, Iain Colin Prentice

## Abstract

The global carbon and water cycles are governed by the coupling of CO_2_ and water vapour exchanges through the leaves of terrestrial plants, controlled by plant adaptations to balance carbon gains and hydraulic risks. We introduce a trait-based optimality theory that unifies the treatment of stomatal responses and biochemical acclimation of plants to environments changing on multiple timescales. Tested with experimental data from 18 species, our model successfully predicts the simultaneous decline in carbon assimilation rate, stomatal conductance, and photosynthetic capacity during progressive soil drought. It also correctly predicts the dependencies of gas exchange on atmospheric vapour pressure deficit, temperature, and CO_2_. Model predictions are also consistent with widely observed empirical patterns, such as the distribution of hydraulic strategies. Our unified theory opens new avenues for reliably modelling the interactive effects of drying soil and rising atmospheric CO_2_ on global photosynthesis and transpiration.

## 1 Introduction

The fundamental dilemma of plants following the C3 photosynthetic pathway is that when stomata, i.e., the tiny ‘valves’ on the surface of leaves, are opened to take in carbon dioxide (CO_2_) for carbon assimilation, water is lost through them via transpiration (Raschke et al., 1976). The plant’s transpiration stream is maintained by negative water potentials (suction pressures) in transport vessels and leaves. Withstanding negative water potentials requires adapted stem and leaf tissues or energy-intensive repair efforts, and extreme water potentials in the xylem can lead to hydraulic failure (Brodribb and Cochard, 2009; Brodribb et al., 2010; Choat et al., 2018). The risks of hydraulic failure increase when water availability declines across the plants’ rooting zones or vapour pressure deficit increases at their leaf surfaces. Plants can avoid hydraulic failure by closing their stomatal openings in response to dry soil and atmospheric conditions. However, closing the stomata also leads to a decline in carbon assimilation, creating a tight coupling between carbon uptake and water loss. At the ecosystem level, this coupling of the carbon and water cycles affects the rates of gross primary production (GPP) and evapotranspiration in response to water stress. Rising atmospheric CO_2_ and precipitation tend to favour increased tree growth rates (Keeling et al., 2017; Guerrieri et al., 2019), whereas rising frequency and intensity of droughts is leading to increased mortality rates (McDowell and Allen, 2015). It has been argued that the interactive effect of saturating growth rates and rising mortality rates is negatively affecting the carbon sink of global forest ecosystems (Brienen et al., 2015). Accurate predictions of the carbon and water cycles thus requires vegetation models which explicitly resolve plant hydraulics (McDowell et al., 2020).

A plant’s hydraulic machinery places key constraints on how much water it can transpire and, consequently, on its stomatal conductance. Considerable effort has gone into the development of stomatal control models with an explicit treatment of plant hydraulics (see reviews (Damour et al., 2010; Y. Wang et al., 2020)). Hydraulically explicit stomatal models have shown success in simulating short-term stomatal responses to drying soil and air on sub-daily and daily timescales (Anderegg et al., 2018; Venturas et al., 2018; Sabot et al., 2020; Eller et al., 2020) and are now being implemented in Earth System Models (Hickler et al., 2006; Bonan et al., 2014; Christoffersen et al., 2016; Kennedy et al., 2019). However, we still lack the understanding of how plant physiology acclimates to the development of soil-moisture drought on daily to weekly timescales and how such longer-term acclimation in turn affects stomatal sensitivity to short-term water stress. Such understanding is especially crucial for predicting stomatal and biochemical responses to novel environments, and for explaining widely observed patterns related to plant hydraulic strategies (Box 1) in a parsimonious way.

### Box 1. Widely observed empirical patterns in plant photosynthetic responses and hydraulic strategies, as targets for model-based predictions.

**Stomatal and biochemical responses to soil and atmospheric drought**

1. As soil moisture decreases, the initial response of plants is to reduce their stomatal opening to alleviate water stress. Since carbon uptake and water loss occur through stomata, photosynthesis and transpiration both decline with stomatal closure, and thus, with decreasing soil moisture (Stocker et al., 2018).
2. As assimilation declines, maintaining photosynthetic capacity becomes increasingly unprofitable. Therefore, in the short term, plant photosynthetic capacity also declines with decreasing soil moisture (Kanechi et al., 1996; Salmon et al., 2020). Yet, in the long term, plants acclimate by shedding their leaves, which reduces transpiration demand and allows assimilation to recover. Thus, in the long term, high photosynthetic capacity can be maintained at the expense of a reduced foliage surface area (Zhou et al., 2016).
3. As vapour pressure deficit increases, the leaf-internal-to-external CO_2_ ratio (*χ*) declines. Various functional forms have been used to describe this decline, which are mostly derived from limited empirical data (Leuning, 1995). A widely used relationship, predicted by simple stomatal optimization models (Medlyn et al., 2011; Prentice et al., 2014), is 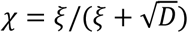, where *D* is vapour pressure deficit expressed as a fraction of atmospheric pressure *ξ* and is a constant. This implies that logit(*χ*) varies linearly with log(*D*) with a slope of −0.5, a value often targeted by modellers (Wolf et al., 2016; Wang et al., 2017). However, analysing data from hundreds of species along aridity gradients, Dong et al. (2020) (Dong et al., 2020) have reported slope values of −0.76 ± 0.15, with remarkable consistency across species. The extent and consistency of these observations suggest that this value ought to be taken as a new target for model-based predictions. **Hydraulic strategies and trait-adaptations**
4. As soil moisture decreases, xylem water potentials become increasingly negative. Extremely negative water potentials create embolisms in the xylem, which have been linked to increased risks of plant mortality (‘hydraulic failure’). To avoid these risks, plants close their stomata before the onset of substantial xylem embolism (Brodribb et al., 2003; Martin-StPaul et al., 2017; Scoffoni et al., 2017; Choat et al., 2018). At the same time, to maximize carbon assimilation, plants tend to keep their stomata open for as long as possible, often close to the point of hydraulic failure. Thus, plants across species operate at extremely low hydraulic safety margins (Choat et al., 2012).
5. Plant traits vary across a continuum of stomatal regulation strategies (Klein, 2014; Papastefanou et al., 2020). At one end are isohydric (drought-avoiding) species that maintain a constant leaf water potential by closing the stomata as soil water potential decreases, at the cost of reduced carbon assimilation. At the other end are extreme anisohydric (drought-tolerating) species that keep their stomata open even in the face of decreasing soil water potential to maintain high CO_2_ uptake, at the risk of hydraulic failure. In between are isohydrodynamic species that maintain a relatively constant soil-to-leaf water potential difference.

The classic stomatal-optimization model by Cowan and Farquhar (1977)(Cowan and Farquhar, 1977) states that plants adjust their stomatal conductance to maximize the total carbon intake gained for a fixed amount of water loss, by assuming a constant unit cost for transpired water. This model implies that plants can save water for future use. However, recent stomatal models recognise that plants competitively consume available water (Wolf et al., 2016). Therefore, an alternative approach conceives the costs of transpiration as arising from the risks of hydraulic failure and the structural and energetic expenditures for withstanding high suction pressures. Thus, many extensions of this classic model explicitly represent plant hydraulics and the associated costs (Wolf et al., 2016; Sperry et al., 2017; Bartlett et al., 2019). These models require an a priori specification of photosynthetic capacity, which then becomes an additional parameter to be fitted to enable accurate predictions of assimilation rates. By contrast, the least-cost optimization framework of Prentice et al. (2014)(Prentice et al., 2014) includes the costs of maintaining carboxylation capacity, reflecting a trade-off between investing in photosynthetic and hydraulic capacities (Wright et al., 2003). Building upon this approach, Wang et al (2017)(Wang et al., 2017) predict acclimated carboxylation capacity using the photosynthetic-coordination hypothesis (Maire et al., 2012; Wang et al., 2017). They further explicitly optimize electron-transport capacity (albeit using a separate optimization criterion) and have been successful in predicting CO_2_ assimilation rates and leaf-internal CO_2_ concentrations across climatic gradients. However, their model requires an empirical factor to account for effects of soil moisture (Stocker et al., 2018, 2020; Lavergne et al., 2020).

Here, we develop a unified first-principles theory combining the photosynthetic-coordination hypothesis with the principles of plant hydraulics within a single optimality framework. Our resulting model simultaneously predicts the stomatal responses and biochemical acclimation (non-stomatal responses) of plants to environments changing on multiple timescales. We test the predictions of our model with published data from soil drought experiments conducted with 18 plant species spanning diverse plant functional types. We show that – with just two parameters – our model correctly predicts key observations related to plant photosynthetic responses and hydraulic strategies, as described in Box 1.

## 2 Model summary

We now list the principles and hypotheses underlying our model in general terms, followed by a summary of the optimality framework, plant traits used in our model, the interpretation of model parameters, and our strategy for testing the model with experimental data. A detailed model description is presented in Methods, and a full derivation of the model is presented in SI Section 1.

### (1) Water-balance principle

Any plant must maintain a continuous transpiration stream across its entire hydraulic pathway (from roots through stem and leaves) to prevent xylem embolism and leaf desiccation (Fig. 1A). Therefore, for a given atmospheric water vapour pressure deficit (VPD), plants adjust their stomatal conductance *g*_s_ such that the atmospheric demand for transpiration is matched by the supply of water from the soil (Sperry and Love, 2015). Since the supply is dependent on the soil-to-leaf water-potential difference Δ*ψ* and the hydraulic properties of the transpiration pathway, this principle predicts Δ*ψ* as a function of *g*_s_ and is widely used in stomatal models that explicitly represent water transport. We use the term ‘principle’ rather than ‘hypothesis’ for this assumption to indicate its rooting in basic physical laws.

**Fig. 1.**
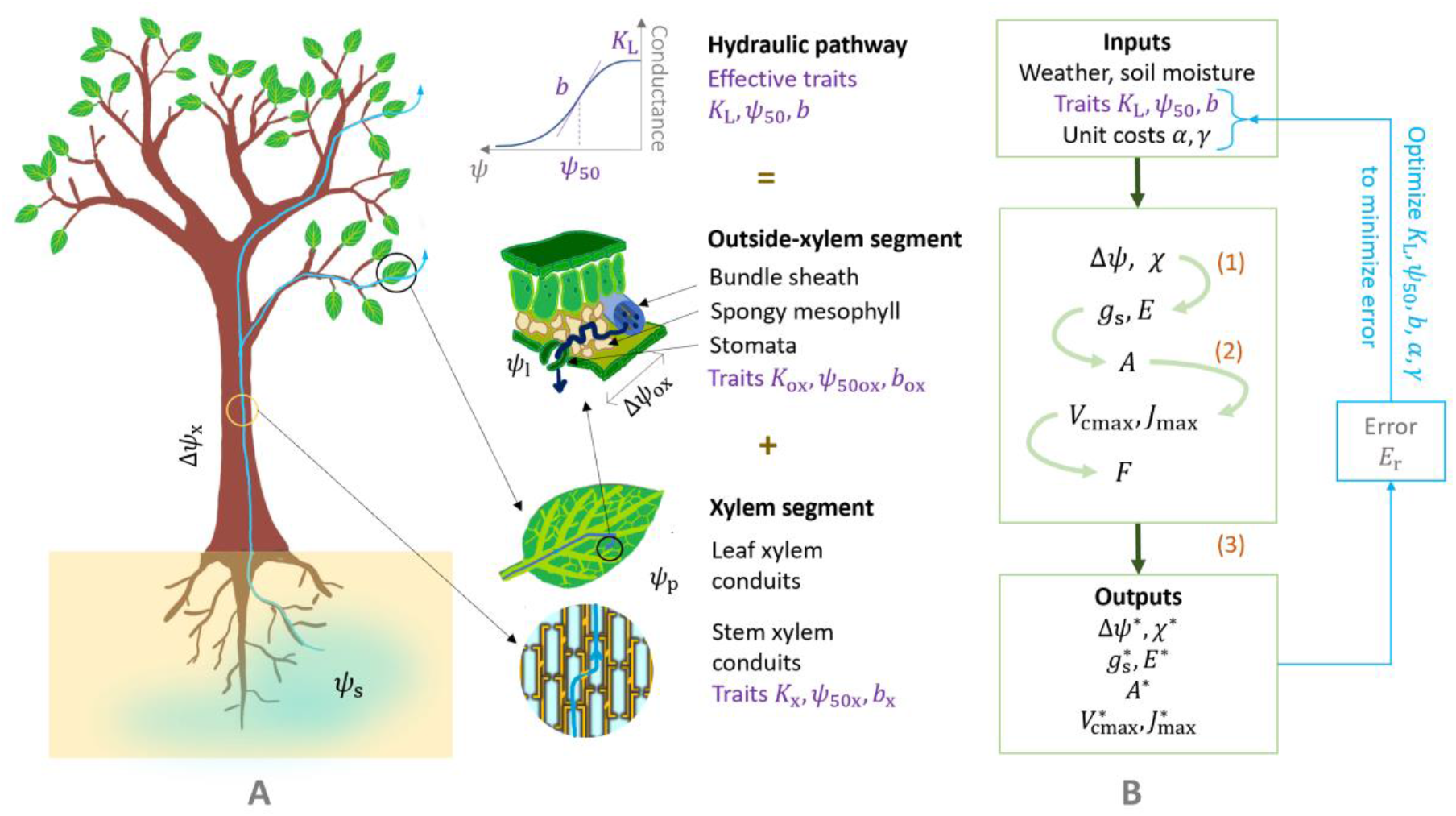
Schematic representation of our model, underlying first principles, and notation. (A) Water-transport pathway. Purple labels indicate the three hydraulic traits that determine the conductance to water flow in the plant. Water potentials are shown at various points along the pathway, *ψ*_s_ in soil, *ψ*_p_ at the end of the xylem segment, *ψ*_l_ and at the end of the outside-xylem segment, i.e., near the stomata. The soil-to-leaf water potential difference *Δψ* =*ψ*_s_ – *ψ*_l_ thus comprises the successive pressure drops along the different segments, i.e., *Δψ*_x_ = *ψ*_s_ – *ψ*_p_ and *Δψ*_ox_ = *ψ*_p_ – *ψ*_l_ along the xylem and outside-xylem segments, respectively. (B) Model-calibration pathway. The model takes as inputs three hydraulic traits (*K*_L_, *ψ*_50_, and *b*) together with two cost parameters (the unit costs of photosynthetic and hydraulic capacities, *α* and *γ*, respectively). It predicts as outputs the optimal values (denoted by asterisks) of stomatal conductance 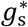, assimilation rate *A*\*, transpiration *E*\*, acclimated photosynthetic capacities 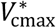 and 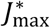, soil-to-leaf water potential difference *Δψ*\*, and leaf internal-to-external CO_2_ ratio *χ*\*. Each variable is first calculated as a function of *Δψ* and *χ*, as shown by the four light-green arrows, from which the optimal combination (*Δψ*\*, *χ*\*) is then calculated by maximizing profit *F* according to Eq. 1. Blue arrows and boxes indicate the process through which the best-fit traits and unit costs for each species are calculated by minimizing the model error. Orange labels indicate the three principles and hypotheses underlying the model, shown next to the processes they affect.

### (2) Photosynthetic-coordination hypothesis

Photosynthetic carbon assimilation is limited by a plant’s capacity for carboxylation *V*_cmax_ and by light availability *I*_abs_, which, together with the electron-transport capacity *J*_max_, determine the rates of biochemical and photochemical reactions governing CO_2_ fixation (Farquhar et al., 1980). In general, the rate of photosynthesis is the minimum of the carboxylation-limited rate *A*_c_ and the light-limited rate *A*_j_. The light-limited rate is further modulated by *J*_max_. Since the carboxylation and electron-transport capacities are costly to maintain, they are hypothesized to acclimate to typical daytime conditions on a weekly timescale, such that the two photosynthetic rates are coordinated, i.e., *A*_c_ = *A*_j_ (Chen et al., 1993; Maire et al., 2012).

### (3) Profit-maximization hypothesis

We posit that, on a weekly timescale (medium-term responses), plants simultaneously optimize their photosynthetic capacity and stomatal conductance to maximize net assimilation (profit, *F*), after accounting for the costs of maintaining photosynthetic capacity and the hydraulic pathway, including the risks of hydraulic failure. On a daily timescale (short-term responses), the acclimated photosynthetic capacities are fixed and plants can optimize only their stomatal conductance. The parameters scaling the photosynthetic and hydraulic costs, *α* and *γ*, respectively, are the only two latent (i.e., not directly observable) parameters in our model and are henceforth called ‘unit costs’.

#### Medium-term responses

To predict the acclimation of photosynthetic capacity on a weekly timescale, we assume that plants independently control their weekly-average stomatal conductance *g*_s_ and their electron-transport capacity *J*_max_ to maximize their net profit *F*, as defined below. After expressing all quantities in Fig. 1B in dependence on *g*_s_ and *J*_max_, or equivalently, in a mathematically more convenient form, in terms of the leaf internal-to-external CO_2_ ratio *χ* and the soil-leaf water potential difference Δ*ψ* (Methods and SI Section 1.3), *F* can be written as

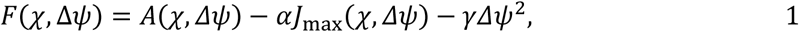

where *A* is the assimilation rate calculated by combining the standard biochemical model of photosynthesis(Farquhar et al., 1980) with the photosynthetic-coordination hypothesis (Eq. 6 in Methods). We find the optimal solution (*χ*\*, Δ*ψ*\*) semi-analytically by first calculating the derivatives of *F* with respect to *χ* and Δ*ψ* analytically (Eq. S14) and then determining their roots numerically.

#### Short-term responses

To predict stomatal responses on hourly and daily timescales, we follow a two-step procedure. First, we find the acclimated photosynthetic capacities using the multivariate optimization described above, driven by a seven-day rolling mean of the soil water potential. Once the acclimated *J*_max_ and *V*_cmax_ are known, *A, g*_s_, and *χ* can all be expressed in terms of Δ*ψ* alone. We again use the net profit in Eq. 1 to optimize Δ*ψ*. In this case, we determine *A* as the minimum of the carboxylation-limited rate *A*_c_ and the light-limited rate *A*_j_. Also, since *J*_max_ is fixed, the term *αJ*_max_ becomes constant and can thus be ignored during optimization.

#### Hydraulic traits

Inside the plant, water first flows through the roots, then through the stem xylem, and then through xylem in leaf veins. After exiting the veins, it flows outside the xylem through the bundle sheath and spongy mesophyll cells, until it evaporates from the stomatal cell walls and diffuses out to the ambient air (Buckley et al., 2015) (Fig. 1A). Under high suction pressure, leaf tissues may desiccate, or xylem vessels may cavitate, causing a decline in plant conductance. Therefore, the flow of water through the plant can be characterized by three effective plant hydraulic traits: (i) the maximum plant conductance per unit leaf area, i.e., the maximum leaf-specific plant conductance *K*_L_, (ii) the water potential *ψ*_50_ that causes a 50% loss of conductance, and (iii) a shape parameter *b* that determines the sensitivity of conductance loss to water potential. There is increasing evidence that the outside-xylem segment of the hydraulic pathway (i.e., the leaf) is the most resistive part of the hydraulic pathway, and thus forms the hydraulic bottleneck of a plant (Sack and Holbrook, 2006; Scoffoni et al., 2017). Therefore, for many species, whole-plant effective hydraulic traits would correspond to the leaf.

#### Interpretation of costs

The photosynthetic costs consist of the costs incurred by maintaining photosynthetic capacities, including the regeneration of RuBP. Since the two photosynthetic capacities are coordinated, these costs are assumed to be proportional to *J*_max_. The hydraulic costs include (i) the construction and respiration costs of the stem and leaves, (ii) the costs of maintaining osmotic potential in the leaves, and (iii) the prospective costs of hydraulic failure. Since these costs are difficult to quantify through mechanistic arguments, we have taken a phenomenological approach and used the expression Δ*ψ*^2^ after assessing several alternative cost expressions including *ψ*_l_, Δ*ψ*, Δ*ψ*^2^, and PLC (percent loss of hydraulic conductivity). A cost expression that is quadratic in Δ*ψ* has also been adopted previously (Wolf et al., 2016).

#### Testing the model with data

We use published data from experiments on 18 species (Table 1) (Zhou et al., 2013), in which plants were grown in greenhouses under controlled conditions and subjected to progressive soil drought; values of *A* and *g*_s_ (and sometimes also of Δ*ψ*) were reported for different values of predawn leaf water potentials, which are indicative of the soil water potential in the plant’s rooting zone. In some experiments, each value of soil water potential was maintained for a long duration, so that photosynthetic capacity could acclimate (species with drought duration = ∞ in Table 1). For such species, we use the multivariate optimization model as described above (Eq. 1). In other experiments, the progression of drought occurred at a natural rate, ranging from 12-60 days (Table 1). For such species, we use the two-step procedure outlined above to obtain the instantaneous values of the assimilation rate and stomatal conductance.

**Table 1.**
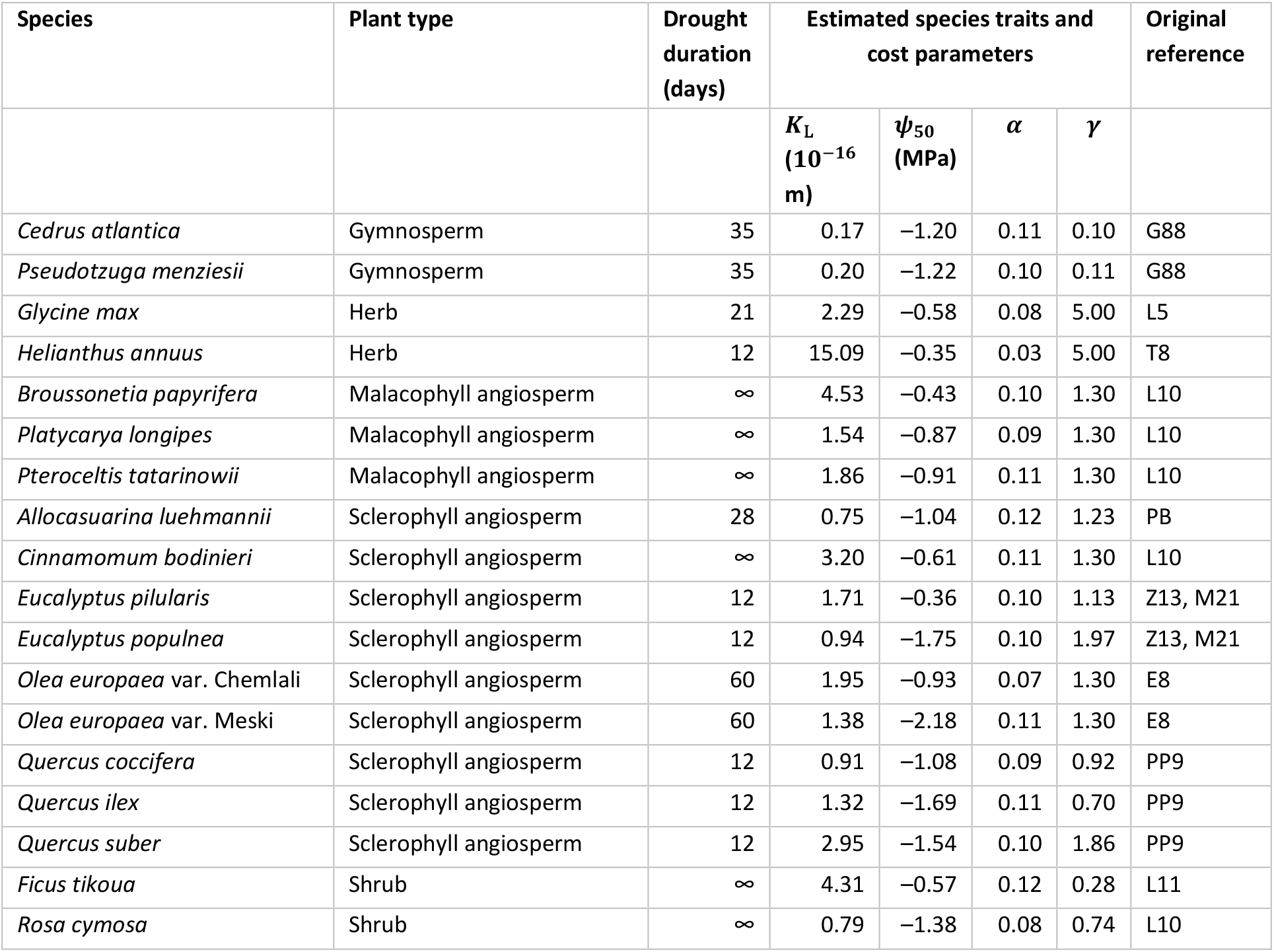
List of species used for testing our model along with their model-estimated trait values. For each species, data on gas exchange for different values of predawn leaf water potential are obtained from Zhou et al. (2013), who originally compiled them from the sources listed in this table. Three hydraulic traits and two cost parameters are estimated using this data. A value of *∞* for drought duration means that soil water potential was maintained at each value for a long time during the experiment, allowing sufficient time for acclimation. To reduce the degrees of freedom in parameter estimation, we set *b* = 1 for all species except *Helianthus annuus*, for which *b* was estimated to be 1.4. G88 - (Grieu et al., 1988) PB - (Posch and Bennett, 2009) L5 - (Liu et al., 2005) L10 - (Liu et al., 2010) L11 - (Liu et al., 2011) E8 - (Ennajeh et al., 2008) Z13 - (Zhou et al., 2013) M21 - (Jiang et al., 2021) PP9 - (Peguero-Pina et al., 2009) T8 - (Tezara et al., 2008)

## 3 Results

### 3.1 Our model correctly predicts the simultaneous decline in gas exchange and photosynthetic capacity under progressive drought

Across 18 species, our model correctly predicts the variation in assimilation rate (*A*), stomatal conductance (*g*_s_), leaf-internal-to-external CO_2_ ratio (*c*_i_: *c*_a_, or *χ*), and soil-to-leaf water potential difference (Δ*ψ*) in response to soil-moisture availability (*ψ*_s_; Fig. 2). Specifically, the shapes of these dependencies closely resemble those observed during experimental drought: Fig. 3 shows predicted and observed responses for two Eucalyptus species from contrasting habitats, and Fig. S1 shows the corresponding responses for all 18 species. Moreover, cross-validation analysis shows that our model generalizes to out-of-sample soil-moisture conditions (Table **S1**).

**Fig. 2.**
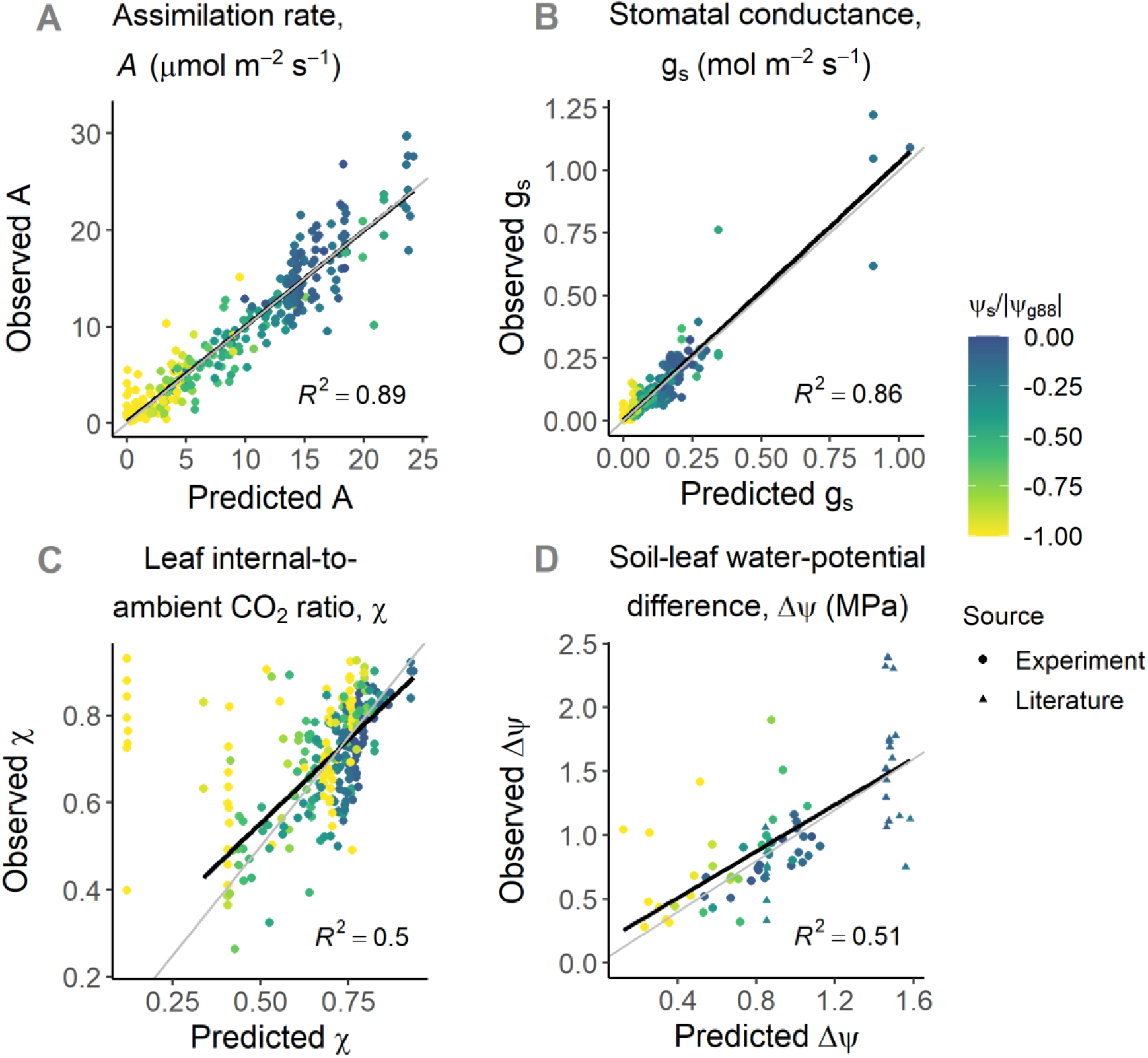
Predicted gas-exchange rates and water relations closely match observations for 18 species. (A-C) Pooled data from all 18 species comparing assimilation rate X, stomatal conductance *g*_s_, and leaf-internal-to-external CO_2_ ratio *χ*, for different values of soil (predawn leaf) water potential *ψ*_s_. (D) Predicted values of soil-to-leaf water potential difference Δ*ψ* compared with observations for (i) six species for which midday leaf water potentials were reported in the corresponding experiments, and thus measured under the same environmental conditions as the gas-exchange rates (circles), and (ii) two species (*Pseudotzuga menziesii* and *Olea europea* var. Meski) for which values were obtained from the literature (Papastefanou et al. 2020) (triangles). Colours indicate soil water potential relative to the stomatal closure point (*ψ*_g88_) of the species; thus, yellow points represent soil water potentials at or beyond stomatal closure. Black lines show linear regressions, while grey lines are the 1:1 lines that represent perfect predictions. In panel (C), we ignore points with *ψ*_s_ < *ψ*_g88_ (yellow points) while calculating the regression line, since there is a known bias in predictions of *χ* beyond stomatal closure (Discussion).

**Fig. 3.**
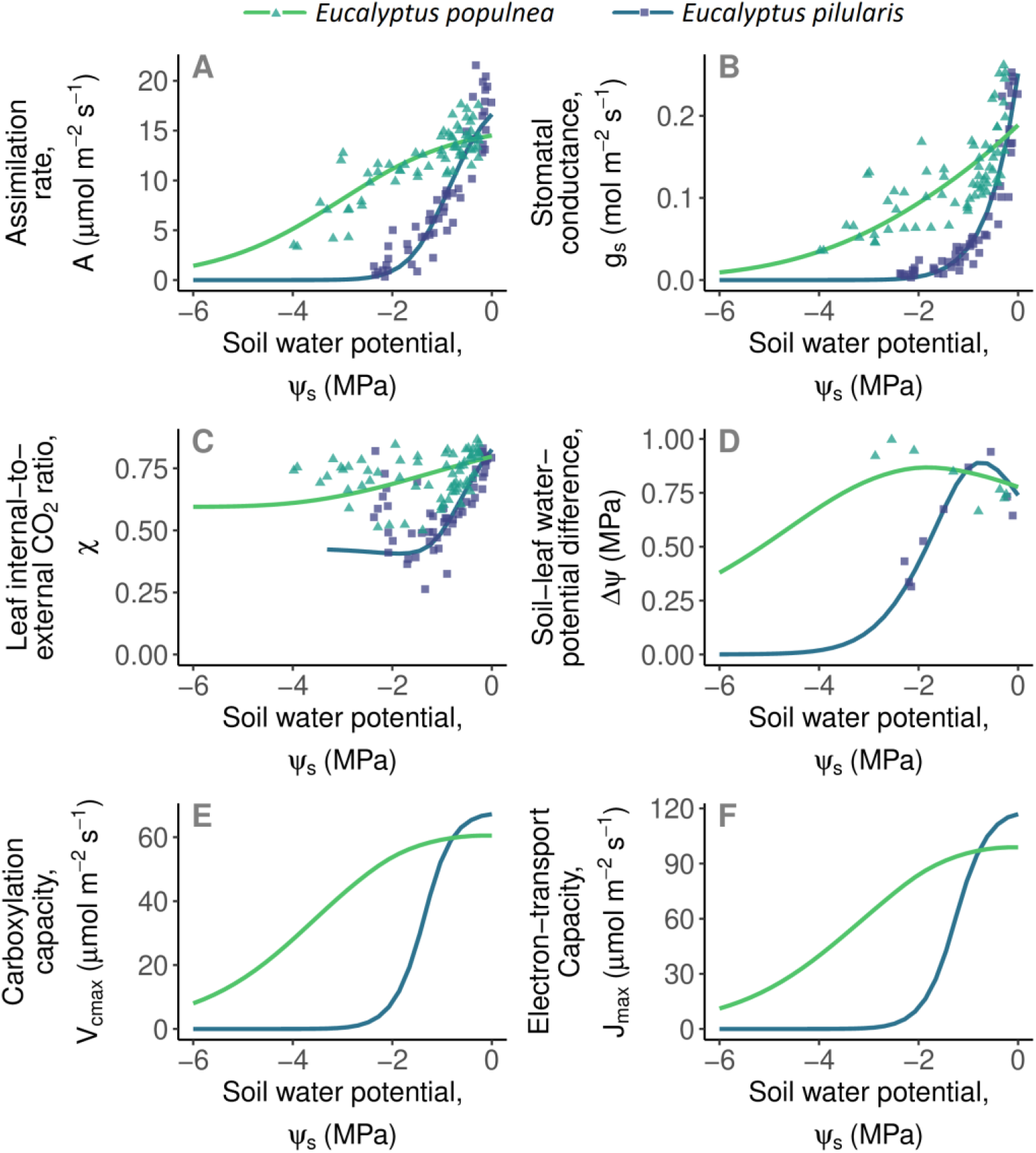
Predicted photosynthetic responses to progressive soil drought closely match observations. Matches are shown here for two *Eucalyptus* species from contrasting climates, and corresponding matches for all 18 species are shown in Fig. S1. Panels show the predicted responses (lines) and observed responses (points) to decreasing soil water potential (*ψ*_s_, measured as pre-dawn leaf water potential): (A) assimilation rate *A*, (B) stomatal conductance *g*_s_, (C) leaf-internal-to-external CO_2_ ratio *χ*, (D) soil-to-leaf water potential difference Δ*ψ*, (E) carboxylation capacity *V*_cmax_, and (F) electron transport capacity *J*_max_. *Eucalyptus pilularis* (blue lines and squares) typically occupies warm and humid coastal areas in eastern Australia, whereas *Eucalyptus populnea* (green lines and triangles) typically occupies semi-arid interior regions of eastern Australia. Since both species were grown in the same greenhouse during the experiment, their contrasting responses reveal genetic adaptations to their native environments. For both species, progressive drought was experimentally induced over 12 days, resulting in a fast instantaneous response of stomatal conductance in combination with a slow acclimating response of photosynthetic capacity. Our model predictions readily account for both responses.

Empirical studies report that photosynthetic capacity (*V*_cmax_ and *J*_max_) declines in response to developing soil drought (Kanechi et al., 1996; Salmon et al., 2020). A unique feature of our model is its ability to predict these responses correctly, qualitatively in accordance with these studies (Fig. 3E-F). Since *χ* depends on both *J*_max_ and *g*_s_, correct predictions of *χ* require predicting both quantities correctly. Therefore, a close match between predicted and observed values of *χ* (Fig. 2C) provides a further quantitative validation of the photosynthetic capacity predicted by our model.

### 3.2 Our model correctly predicts the photosynthetic responses to vapour pressure deficit and other atmospheric variables

Our work builds on the principles introduced by Wang et al. (2017) (Wang et al., 2017), and thus inherits their capacity to accurately predict (Lavergne et al., 2020) photosynthetic responses to temperature, atmospheric CO_2_, and light intensity (Fig. S2C-E). Furthermore, by explicitly modelling leaf hydraulics, our model delivers improved predictions of photosynthetic responses to soil moisture and vapour pressure deficit (Fig. S2A-B).

Furthermore, the functional relationship between logit(*χ*) and log(*D*) predicted by our model shows a close match with observations (Box 1, point 3). In particular, our model predicts this relationship to be linear with slope values of −0.695 ± 0.015 (Fig. 4A). These predicted slope values are well within the confidence interval reported by Dong et al. (2020) (Dong et al., 2020). Also, we find that this slope is negatively correlated with *ψ*_50_ (Fig. 4B), such that species with highly negative *ψ*_50_ have less negative slope values. Since earlier datasets were dominated by temperate evergreen species, this could explain why a slope value of −0.5 predicted by previous models was supported by such datasets. We offer this predicted correlation between the slope and *ψ*_50_ as an empirically testable prediction for future studies.

**Fig. 4.**
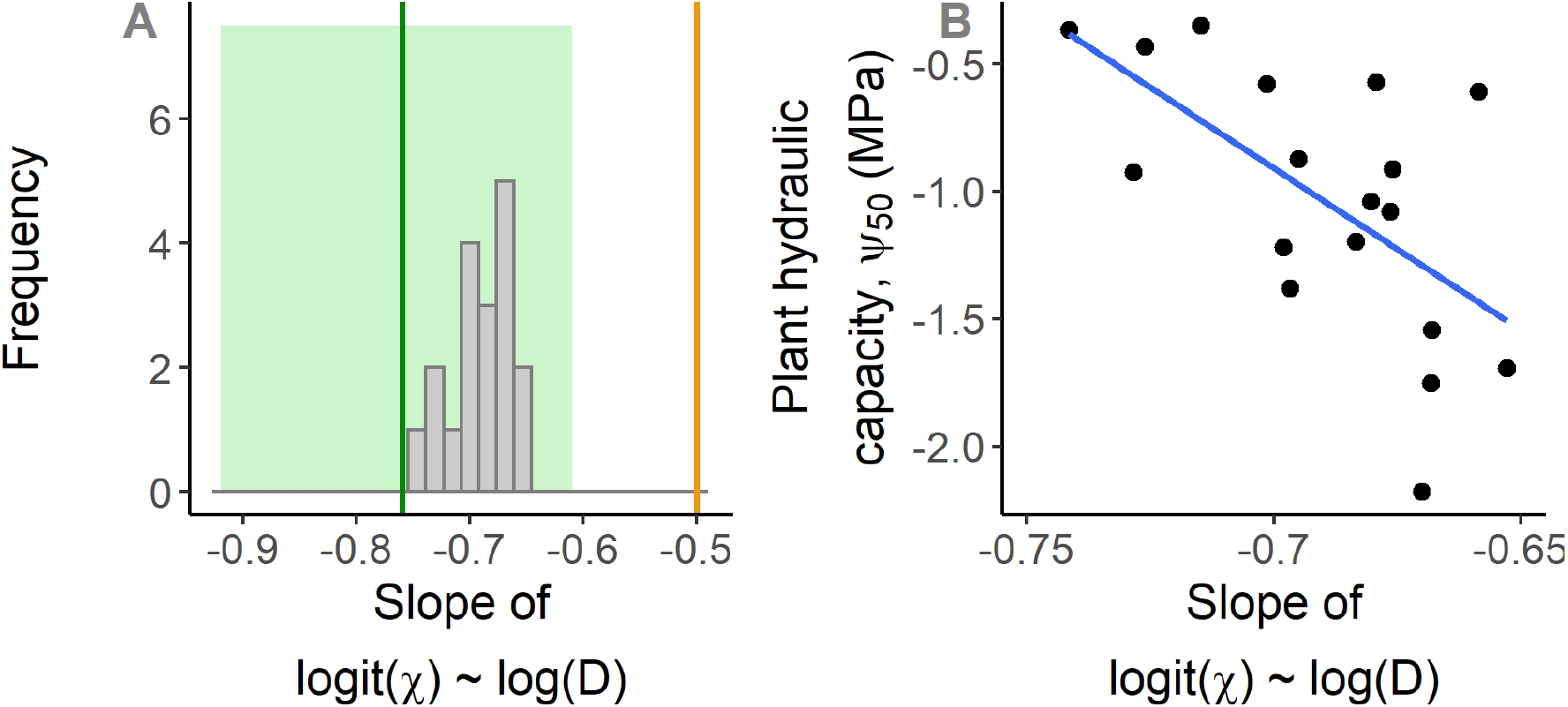
Our model correctly predicts the response of *χ* to vapour pressure deficit. (A) The model-predicted distribution of the slope of the relationship between logit(*χ*) and log(D) for the analysed species (grey bars) is well within the range reported by Dong et al (2020) (their reported mean and confidence interval is shown by the green line and green region, respectively). It is significantly different from −0.5 (orange line; a one-sample t-test shows a predicted mean slope of −0.695 and a 95% confidence interval of [−0.71, −0.68]). For each species, we calculate the predicted slope by varying vapour pressure deficit in the range 5–5000 Pa while keeping other environmental parameters constant (at values reported in the respective experiments, with *ψ*_s_ = 0) and using fitted trait values (Table 1). (B) This slope is correlated with the *ψ*_50_, with more negative slopes observed for species with less negative *ψ*_50_ (drought avoiders). This could be a reason why earlier datasets supported a slope value of −0.5, as such datasets were often dominated by temperate evergreen species, which are typically characterised by highly negative values of *ψ*_50_.

### 3.3 Model-predicted patterns of hydraulic strategies are consistent with widely reported empirical observations

In this section, we compare several widely observed empirical patterns among plant hydraulic traits with the corresponding model-predicted patterns. This qualitative comparison allows us to validate our model at an even deeper level.

First, we compare distribution of the degree of anisohydricity (Box 1) observed in our model to that reported in the empirical literature (Martínez-Vilalta et al., 2014). For each species, the degree of anisohydricity is determined by the slope of the relationship between the water potential in the leaf (*ψ*_l_) and in the soil (*ψ*_s_) measured at low *ψ*_s_ (slope <1 for isohydric, =1 for isohydrodynamic, and >1 for anisohydric species). The observed global distribution of these slopes peaks at approximately 1, suggesting that the global majority of species follow the isohydrodynamic strategy (Martínez-Vilalta et al., 2014). The corresponding distribution predicted by our model lies within the observed distribution (Fig. 5A). Similarly, the predicted distribution of typical operating water potentials (*ψ*_l_ at *ψ*_s_ = 0) also closely matches the corresponding empirically observed distribution (Fig. S3).

**Fig. 5.**
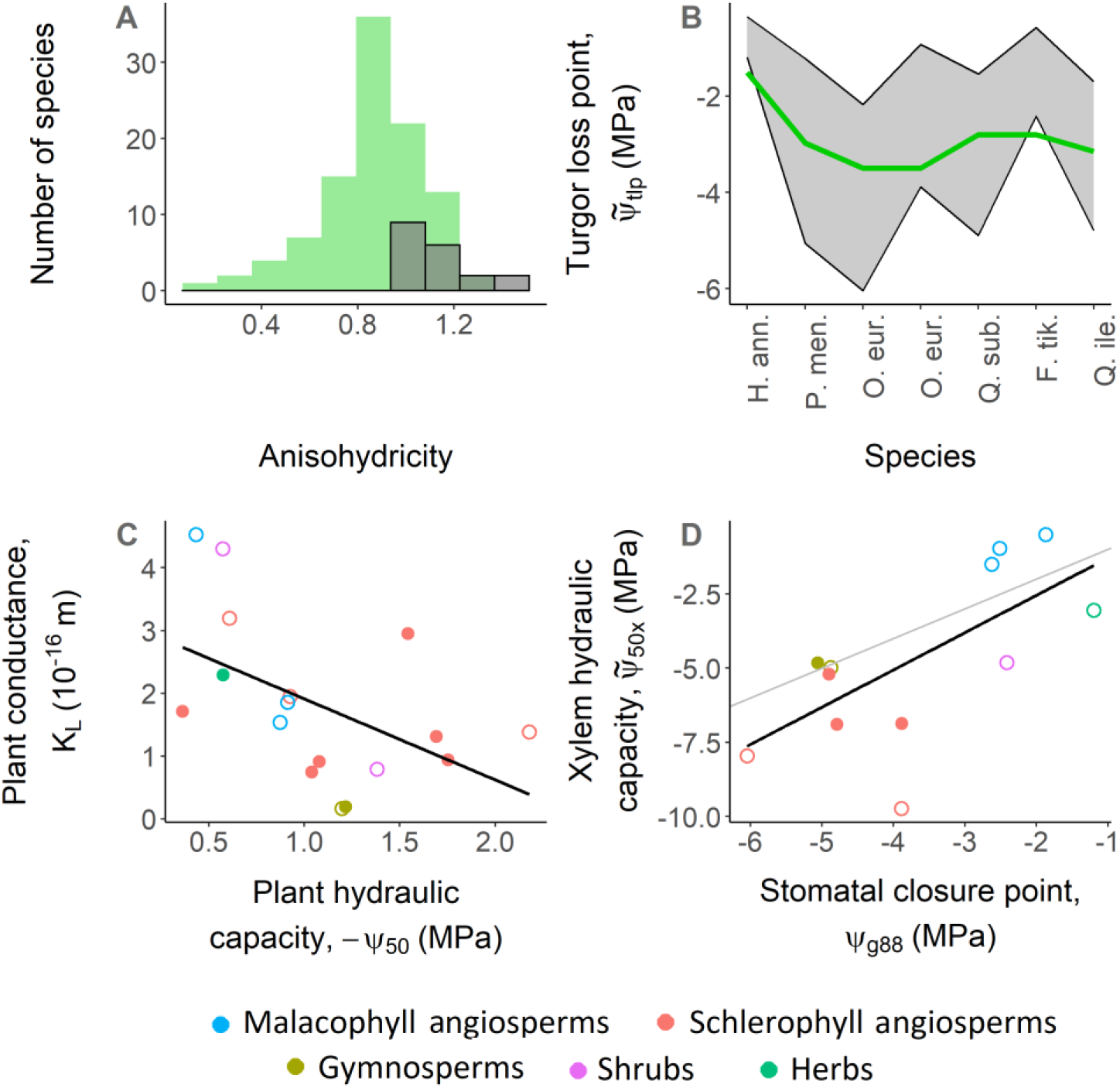
Our model predictions are consistent with widely observed empirical patterns. (A) The predicted distribution of the degree of anisohydricity among the 18 analysed species (grey bars) lies within the observed global distribution (green bars; as reported in Martinez-Vilalta et al (2014)). (B) Consistent with empirical observations, the turgor loss point (thick green line) lies between the water potential at 50% loss of plant conductivity (*ψ*_50_; top black line) and the water potential at 88% stomatal closure (*ψ*_g88_; bottom black line). (C) Plant conductance (*K*_L_) is weakly negatively correlated with *ψ*_50_, with no species having high values of both traits, implying a weak safety-efficiency trade-off in line with empirical observations. (D) When leaf water potential is at *ψ*_g88_, the loss of xylem conductivity is typically less than 50% (implied by observed xylem hydraulic capacity 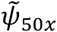 being less than model-predicted *ψ*_g88_), which means that plants close their stomata before the onset of substantial xylem embolism. Furthermore, the difference between the regression line (black) and the 1:1 line (grey) is low, implying that the hydraulic safety margin 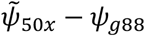 is small on average. Sources for values of 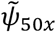 are given in Table S2. Closed circles indicate species for which *γ* was estimated using data on Δ*ψ*, whereas open circles refer to species for which such data was not available and for which we therefore used an average value of *γ* estimated for the respective plant types.

Second, we compare the empirical and model-predicted relationships between leaf turgor loss point and leaf hydraulic capacity. Empirical observations show that the turgor loss point lies between the point where the leaf loses 50% conductivity (*ψ*_50_) and the point of stomatal closure (*ψ*_g88_) (Bartlett et al., 2016). Our model predictions are consistent with this observation in all species for which data are available (7 out of the 18 species) (Fig. 5B).

Third, we compare how plant hydraulic capacity (*ψ*_50_), a measure of hydraulic safety, covaries with plant conductance (*K*_L_), a measure of hydraulic efficiency. Global data reveals a trade-off between safety and efficiency, i.e., no plants score high on both, but only a weak correlation between them, i.e., many plants score low on both. Consistent with these observations, we find only a weak correlation between model-estimated values of *ψ*_50_ and *K*_L_, with a few species having low values of both traits, but no species having high values of both (Fig. 5C).

Fourth, we compare how empirical and model-estimated stomatal closure point relates to xylem hydraulic capacity. Empirical observations show that stomatal closure occurs before the onset of substantial xylem embolism (Brodribb et al., 2003; Martin-StPaul et al., 2017; Scoffoni et al., 2017; Skelton et al., 2018), which likely is an adaptation to prevent plant mortality during drought (Choat et al., 2018). At the same time, the minimum water potential experienced by the leaves (*ψ*_min_) is close to the water potential at which the xylem loses 50% conductivity (*ψ*_50x_), leading to extremely low hydraulic safety margins (Choat et al., 2012). Both of these observations are matched by our model, confirmed by a close correlation between the model-predicted values of the stomatal closure point (*ψ*_g88_, which is a proxy of *ψ*_min_) and empirically observed values of *ψ*_50x_ (Fig. 5D).

## 4 Discussion

We have presented an analytical trait-based optimality model, unifying plant photosynthesis and hydraulics to predict the stomatal responses and biochemical acclimation of plants to changing hydroclimates. Tested with experimental data on 18 species, we find that our model correctly predicts the stomatal and photosynthetic responses to soil drought and the dependencies of photosynthesis on vapour pressure deficit, temperature, and CO_2_. We also find that it is consistent with several widely observed empirical patterns. To our knowledge, our multivariate optimization model is the first to optimize photosynthetic capacity concurrently with stomatal conductance, thereby allowing us to predict the observed S-shaped decline of *V*_cmax_ in response to drying soil from first principles. Furthermore, explicit inclusion of plant hydraulics in our model allows for separating the stomatal responses to atmospheric drought and soil-moisture availability, which could be particularly useful for remote-sensing based models of GPP.

### 4.1 Comparison with other stomatal optimization models

Leaf photosynthesis is known to be jointly constrained by stomatal and non-stomatal limitations. A vast majority of photosynthesis models account only for stomatal limitations, where stomatal conductance is optimized to maximize photosynthetic gain. Non-stomatal limitations, such as the constraints imposed by leaf mesophyll, photosynthetic capacity, and sugar transport, have received attention only in the most recent stomatal models. Even such models account for them using a pre-determined functional response, in which mesophyll conductance (Dewar et al., 2018) or photosynthetic capacity (Hölttä et al., 2017; Sperry et al., 2017; Dewar et al., 2018) is scaled in a prescribed way with stomatal conductance. By contrast, our model can account for non-stomatal limitations without the need to specify photosynthetic capacity a priori. This is especially important for out-of-sample predictions, when applying the model to conditions outside the domain of environmental factors used for calibration, such as elevated CO_2_ levels, and when applying it to species for which empirical estimates of photosynthetic capacity are not available.

Almost all current models of stomatal optimization focus on water transport through the xylem. By contrast and in line with growing evidence (Sack and Holbrook, 2006; Scoffoni et al., 2017), we have hypothesized that the leaf is the hydraulic bottleneck of the plant. This implies that the fitted hydraulic traits in our model would correspond to the leaf. Indeed, we have found support for this hypothesis, as fitted values of *ψ*_50_ in our model are much less negative than the corresponding values for xylem. We offer the fitted values of these traits (*ψ*_50_ and *K*_L_) as testable predictions that can be validated by explicitly measuring these traits on the leaves of the corresponding species.

### 4.2 Model assumptions and extensions

Under extreme hydroclimatic conditions, such as extremely dry or flooded soils, or extremely low atmospheric CO_2_ levels, our model predictions may deviate slightly from observations. For example, most species in our dataset show an increase in the leaf-internal-to-external CO_2_ ratio (*χ*) after stomatal closure (Fig. 3). The model-predicted *χ* does not increase, asymptotically approaching a constant value instead (Fig. 3, Eq. S16). The increase in *χ* is due to a build-up of CO_2_ in the leaf, which can happen via two mechanisms: (i) if dark respiration continues even after assimilation has ceased, or (ii) if assimilation and respiration decline together due to reduced photosynthetic activity while CO_2_ continues to ‘leak in’ through the leaf cuticle (Boyer et al., 1997). Future research could identify which of these is observed in leaves: since the source of CO_2_ is different in the two mechanisms (plant metabolism or ambient air), they can be distinguished based on whether the build-up is detected also in δ^13^C measurements. Our model could be extended to include these mechanisms, but this would not affect the predicted assimilation rates because these mechanisms are relevant only after stomatal closure.

For simplicity, we have assumed an effective hydraulic pathway by representing the pathway’s different segments (leaves, xylem, and roots) by a single effective segment. This assumption is expected to hold for many species in which one segment (typically the leaf) contributes most of the resistance to water flow. However, there could be species in which multiple segments have similar conductivity. The numerical version of our model can readily be extended to account for segmented pathways. In SI Section 2, we present a possible extension of the model considering two segments (xylem and outside-xylem). Such an extension would be useful also for understanding the relative roles of the xylem and leaves in stomatal control and drought survival.

One detail we have not included in our model is the leaf’s energy balance. This was done to avoid making the model too complex and to enable the semi-analytical solution. Nevertheless, under drying soil, reduced transpiration raises the leaf temperature, which in turn affects the temperature-dependent photosynthetic parameters and the dark-respiration rate. Inclusion of this effect in our model could therefore be a promising direction for further research.

Plants respond to increasing water stress on multiple timescales and at multiples scales of organization (leaves, whole plants, and even stands). Our model accounts for leaf-level responses on daily and weekly timescales, capturing the role of stomatal closure in preventing damagingly negative stem water potentials. Our model could be extended to develop optimality-based first-principles models for the plant level (Deans et al., 2020) to account for responses beyond the point of stomatal closure, such as the shedding of leaves on a monthly timescale, either in full to prevent loss of water through cuticular tissue (Choat et al., 2018) or in part to reduce transpiration demand and continue photosynthesis (Zhou et al., 2016). Similarly, changes to the characteristics and architecture of the transpiration pathway, as reflected in stem traits (Rungwattana et al., 2018), could be modelled on yearly to decadal timescales. Trait adaptation on centennial or millennial timescales could be modelled by embedding our leaf-level optimality theory into evolutionary models (Dybzinski et al., 2011; Hikosaka and Anten, 2012; Franklin et al., 2020).

In our model, on the shortest timescale (minutes-days), plants may optimize leaf water potential for a fixed (acclimated) photosynthetic capacity. On the timescale of weeks, plants additionally adjust their photosynthetic capacities. In principle, the weekly timescale can either be modelled with nested optimization (i.e., optimizing daily, or sub-daily, stomatal conductance for a given photosynthetic capacity, and optimizing weekly photosynthetic capacity by maximizing the total profit over a week), or with simultaneous optimization (i.e., optimizing both variables together by assuming a constant environment during the week, representative for mean daytime conditions). In this work, we have taken the latter approach for theoretical and computational simplicity, but the alternative nested approach is worth exploring in future work.

### 4.3 Implications for global vegetation modelling

Taking advantage of the increasing availability of global data on plant traits, our model can be applied at the global scale by making a few additional assumptions. For the species in our dataset, we find that the photosynthetic unit cost *α* lies within a narrow range of values (0.08–0.12), which could therefore be treated as a constant. Furthermore, variations in the hydraulic unit cost *γ* are primarily driven by differences between plant types, with relatively lesser variability within plant types, suggesting that *γ* could also be treated as a constant within plant types (Fig. S4). To infer the global distribution of *α* and *γ*, our model can be used in a Bayesian framework on global data on gas-exchange measurements. When the leaf-level hydraulic traits *ψ*_50_ and *K*_L_ are not available, they can be estimated from other widely measured traits: *ψ*_50_ is correlated with xylem hydraulic capacity (Fig. 5D), and *K* is correlated with specific leaf area (Fig. S4B).

Accurate models of plant photosynthesis are crucial for improving projections of the global carbon and water cycles, because photosynthesis and transpiration by terrestrial plants account for 56% and 30% of the global fluxes of carbon dioxide and water, respectively (Jasechko et al., 2013; Le Quéré et al., 2018). It is especially important to develop models that can generalize to new climatic conditions, because the projected increase in the frequency and intensity of droughts may lead to unprecedented climatic conditions in future. The inclusion of plant hydraulics into vegetation models has been shown to improve predictions of global productivity and evapotranspiration (Hickler et al., 2006; Bonan et al., 2014; Christoffersen et al., 2016; Kennedy et al., 2019; Eller et al., 2020), as well as predictions of the spatiotemporal diversity of vegetation (Xu et al., 2016). Spearheading the development initiated by these studies, our model is ideally suited for being embedded into global vegetation models: by accounting for biochemical acclimation, plant hydraulics, and photosynthetic trade-offs through optimality principles, our model can extend to new species and new environmental conditions with a raised degree of confidence. Furthermore, accounting for photosynthetic and hydraulic costs is bound to yield more accurate estimates of the energy spent on resource acquisition and, consequently, of the resources available for growth and reproduction. Therefore, embedding our model of photosynthesis and hydraulics into a demographic model of plant communities can help improve the scaling of assimilation and transpiration from the leaf level to the whole-plant level, and even from plants to communities, thus paving the way for more accurate and robust land-surface models.

## 5 Methods

Our model consists of three key components or modules, corresponding to the three principles and hypotheses – a water-transport module to account for plant hydraulics and water balance, a photosynthesis module to account for photosynthesis and the photosynthetic-coordination hypothesis, and a profit-maximization module to implement the optimization. Here, we describe the equations used for each module, as well as our strategy for model calibration. Full derivations of the equations can be found in SI Section 1.

### 5.1 Water-transport module

We model water transport using Darcy’s law applied to small cross-sections of the hydraulic pathway (SI Section 1.1.4). In principle, our model of water transport can explicitly represent multiple segments (SI Section 2), but for simplicity, we represent the entire pathway by a single ‘effective segment’ with traits *K*_L_, *ψ*_50_, and *b*. Thus, our hydraulic model is mathematically similar to the one described by Sperry et al. (2017)(Sperry et al., 2017) for xylem water transport, but the effective hydraulic traits in our model correspond not necessarily to the xylem, but to the most resistive segment in the pathway. As the leaf is the hydraulic bottleneck of the plant for many species (Sack and Holbrook, 2006; Scoffoni et al., 2017), the traits modelled here would likely correspond to the leaf in such species. In some other species, the root cortex might be the segment with the fastest loss of conductivity (Sperry et al., 2008). For species in which the different segments may be colimiting, care must be taken when comparing estimated effective traits with observations because empirically observed values may differ depending on which part of the plant (root, stem base, apex, or leaf) was sampled, and corrections may have to be made to account for the effects of vessel tapering.

As leaf tissues desiccate or xylem vessels cavitate under high suction on the water column, the conductivity *κ* of any cross-section of the pathway declines as the water potential becomes increasingly negative. This decline in conductivity is phenomenologically described by a so-called vulnerability curve *P*(*ψ*), such that *κ*(*ψ*) = *κ*(0)*P*(*ψ*). The vulnerability curve is typically described by a Weibull function with two parameters – the water potential *ψ*_50_ at which 50% conductivity is lost and a shape parameter *b* that determines the sensitivity of conductivity loss to water potential,

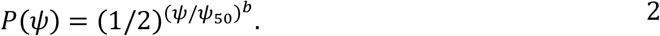

Water potential drops continuously along the hydraulic pathway, from *ψ*_s_ in the soil to *ψ*_l_ at the leaf surface, with a continuous decline in conductivity along the pathway. The volumetric flow of water per unit leaf area in the pathway, *Q*, is therefore described by a differential equation (see SI Section 1.1.4 for derivation), which can be solved for *Q* as follows,

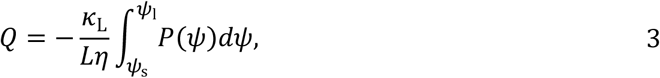

where *κ*_L_ is the conductivity of the pathway per unit leaf area, *L* is the effective length of the pathway, and *η* is the dynamic viscosity of water. The composite term *κ*_L_/*L* = *K*_L_ is the plant conductance per unit leaf area (SI Section 1.1). To keep the number of parameters low, in order to prevent overfitting of the model to data, we use Eq. 3 to model water flow. In SI Section 2, we propose an extension of this model, deriving an expression for *Q* by explicitly accounting for the xylem and outside-xylem segments of the hydraulic pathway, each with its own traits.

The water-balance principle states that the atmospheric demand for water imposed by vapour pressure deficit at the leaf surface matches the supply of water from the soil via the stem and leaf segments of the hydraulic pathway. The transpiration rate at which water vapour diffuses out of the leaf into the atmosphere is given by (see SI Section 1.1.6 for derivation),

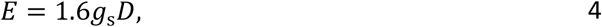

where *g*_s_ is the stomatal conductance and *D* is the atmospheric vapour pressure deficit divided by the atmospheric pressure. This rate *E* equals the rate *Q* at which water enters the leaf according to Eq. 3, which allows us to calculate *g*_s_ by solving

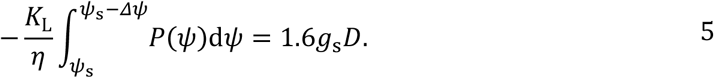

### 5.2 Photosynthesis module

The assimilation rate *A* is calculated from the Farquhar-von Caemmerer-Berry biochemical model (Farquhar et al., 1980) (SI Section 1.2.1), with the photosynthetic capacities *J*_max_ and *V*_cmax_ linked through the photosynthetic-coordination hypothesis (Fig. 1B; SI Section 1.2.4). Temperature responses of photosynthesis parameters, such as the Michaelis-Menten coefficient and the light-compensation point, are modelled with Arrhenius functions for enzymatic rates as described in Stocker et al. (2020)(Stocker et al., 2020).

The photosynthetic-coordination hypothesis states that under typical daytime conditions, assimilation operates at the point of co-limitation, such that the carboxylation-limited and light-limited assimilation rates are equal. With this assumption, the co-limited assimilation rate can be written as (SI Section 1.2)

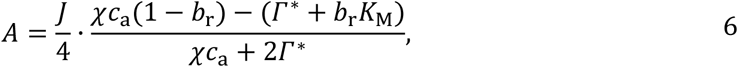

where *J* is the effective electron-transport capacity, which increases with light availability *I*_abs_ but saturates due to limitation by the leaf’s intrinsic electron-transport capacity *J*_max_,

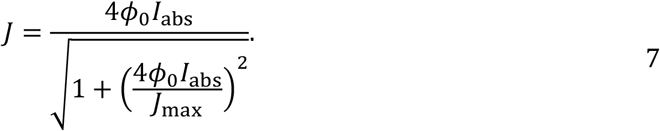

Here *c*_a_ is the atmospheric CO_2_ concentration, *χ* is the ratio of the leaf-internal and external CO_2_ concentrations (*c*_i_: *c*_a_), *Γ*\* is the light compensation point, is the Michaelis-Menten coefficient for C3 photosynthesis, *ϕ*_0_ is the quantum yield efficiency, *I*_abs_ is the absorbed photosynthetically active radiation, and *b*_r_ is the ratio of dark respiration to carboxylation capacity (dark respiration is assumed to be proportional to carboxylation capacity, i.e. *R*_d_ = *b*_r_*V*_cmax_). Temperature dependencies of *Γ*\* and are modelled according to Stocker et al., (2020) (Stocker et al., 2020).

The ratio *b*_r_ also has a weak dependence on temperature (H. Wang et al., 2020), which we have ignored in this work. Variation in *J*_max_ in response to light and water availability (by optimization) implies a coordinated variation in both carboxylation and electron transport capacities.

### 5.3 Profit-maximization module

We assume that plants maximize net assimilation (or profit, *F*) defined in Eq. 1. Without loss of generality, we have assumed that the unit benefit of assimilation is one, i.e., *α* and *γ* represent the ratios of the unit costs to unit benefits of assimilation. To optimize Eq. 1, we express all quantities in terms of the two independent variables *χ* and *Δψ* and set the gradient of the profit function to 0. This can be done analytically (Eq. S14). However, except in the special case of strong *J*_max_ limitation, the roots of the gradient must be found semi-analytically (SI Section 1.3.2). Solving for optimal *χ*\* and *Δψ*\* in turn allows us to predict the optimal photosynthetic capacities (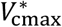 and 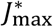), stomatal conductance (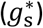), and CO_2_ assimilation rate (*A*\*).

### 5.4 Parameter estimation and model testing

We drive the model with environmental variables (temperature, vapour pressure deficit, light intensity, and CO_2_) as specified in the experimental studies. Other parameters used in the model are as follows: *ϕ*_0_ = 0.087, *b*_r_ = 0.02. In the case of instantaneous responses, we use the daily maximum light intensity under growth conditions to calculate the acclimated response, and saturating light intensity (as specified in the studies) to calculate the instantaneous response, so as to mimic the conditions present during LiCor measurements.

Since measurements of effective whole-plant hydraulic traits are not readily available, we treat them as model parameters and estimate them along with the two cost parameters. Values of Δ*ψ* were reported for 6 of the 18 species from the same drought experiments. We supplement Δ*ψ* observations with typical values reported in the literature (Papastefanou et al., 2020) for two additional species (*Pseudotzuga menziesii* and *Olea europea* var. Meski). For such species (for which measurements of Δ*ψ* are available), we calibrate five parameters (*α, γ, ψ*_50_, *b*, *K*_L_) by minimizing the sum of squared errors (*E*_r_) between predicted and observed values of *A, g*_s_, *χ*, and Δ*ψ*, defined as

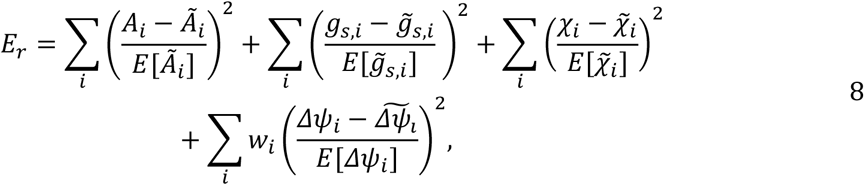

where *i* represents different values of *ψ*_s_, *E*[] denotes the mean value, and variables with tilde (e.g., 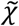) represent observations.

From this calibration, we obtain the mean estimated value of *γ* for each plant type. For all other species (for which measurements of Δ*ψ* are not available), we use this mean value of *γ* and estimate the remaining four parameters. To further reduce the degrees of freedom in model parameterization, we fix the value of *b* = 1 for all species except *Helianthus annuus*, for which *b* also had to be estimated. For each species for which data on Δ*ψ* is available, we evaluate model performance using five-fold cross-validation (or leave-one-out cross-validation when data points are limited).

### 5.5 P-hydro R package

R code to run our model (P-hydro) is provided as an extension of the *rpmodel* package (https://github.com/jaideep777/rpmodel/tree/hydraulics), with options to use the acclimating and instantaneous responses. It also provides an option to use the semi-analytical solution derived in this work, or to directly optimize the profit function numerically. The numerical implementation also allows for quick extension of the model with different profit and cost functions.

## 6 Acknowledgements

We thank Belinda Medlyn and Oskar Franklin for discussions and feedback on the manuscript. JJ and UD acknowledge funding from the European Commission through a Marie Skłodowska-Curie Actions fellowship (Grant No. 841283 – Plant-FATE). JJ, FH, and UD gratefully acknowledge funding from the International Institute for Applied Systems Analysis (IIASA) and the National Member Organizations that support the institute. JJ also gratefully acknowledges support from the Divecha Centre for Climate Change, Indian Institute of Science in the form of initial funding. This work is a contribution to the Imperial College initiative on Grand Challenges in Ecosystems and the Environment and has received funding from the European Research Council (ERC) under the European Union’s Horizon 2020 research and innovation programme (grant agreement No: 787203 REALM). BDS was funded by the Swiss National Science Foundation grant no. PCEFP2_181115.

