## Supplementary information for "Towards a unified theory of plant photosynthesis and hydraulics"

### Supplementary Information:
### Towards a unified theory of plant photosynthesis and hydraulics

Jaideep Joshi, Benjamin D. Stocker, Florian Hofhansl, Shuangxi Zhou,
Ulf Dieckmann, and Iain Colin Prentice

### 1 Model description

Our model comprises three components specified as modules: the water-transport module (Section 1.1), the photosynthesis module (Section 1.2), and the optimization module (Section 1.3). Variables and notations for the three modules are summarized in Tables M1-M3, respectively.

#### 1.1 Water-transport module and water-balance principle

In this section, we introduce the biophysical concepts and terminology necessary for describing plant hydraulics, describe the equation governing water flow within a plant as well as its solution, and specify and apply the water-balance principle.

##### 1.1.1 Water potential

The water potential $\psi$ measures the potential energy of water per unit volume, with the value $\psi = 0$ corresponding to liquid water in fully saturated soil under reference conditions. Accordingly, the water potential $\psi$ becomes increasingly negative as water ascends through a plant's hydraulic pathway, reaching very negative values when it exits the plant into the atmosphere as water vapour through the stomata.

##### 1.1.2 Volumetric water flow

The volumetric flow rate $q$ of water through a pipe is given by Darcy's law,

$$q = \frac{k}{\eta}\Delta\psi,$$

where $k$ is the conductance of the pipe to water flow depending on the pipe's cross-sectional area $A$ and length $L$, $\eta$ is the viscosity of water, and $\Delta\psi$ is the water-potential difference between the beginning and end of the pipe. In plant hydraulics, it is often convenient to express the water flow rate $q$ in terms of a flow rate $Q$ per unit surface area,

$$Q = \frac{q}{A} = \frac{K}{\eta}\Delta\psi,$$

where $K = k/A$ is the conductance of the pipe per unit surface area. Since we follow the widespread convention of expressing $Q$ as the flow rate per unit leaf area rather than per unit sapwood cross-sectional area, we choose to also express the conductance $K$ per unit leaf area.

##### 1.1.3 Hydraulic conductivity

The longer water must travel through a pipe, the more resistance – and thus, less conductance – it experiences: specifically, the conductance $K$ is inversely proportional to the pipe length $L$,

$$K = \frac{\kappa}{L}.$$

where $\kappa$ is the hydraulic conductivity, which measures the permeability of the conducting medium. Since $K$ is already expressed per unit surface area and the inverse relationship with length is accounted for as indicated above, $\kappa$ is independent of all dimensions of the pipe, instead characterising the conducting medium.

In the plant, water flows through the xylem in the stem and in the leaves, and then outside the xylem in the leaves. There are two sources of variation in conductivity along the hydraulic pathway. First, conductivity depends on the internal structure of the xylem and of the leaf mesophyll, such as the number and interconnectivity of xylem conduits and mesophyll cells. This internal structure often changes systematically with plant height, such as through the continuous tapering of the xylem vessels. Therefore, conductivity can be expressed as a function of the distance $h$ from the plant base (Fig. M1). Second, as water potential decreases, individual xylem conduits embolize, leading to a reduction in the xylem conductivity. Similarly, the permeability of the cell membranes in the leaf decreases with increasing water tension, causing a decline in the outside-xylem conductivity. Thus, conductivity is also a function of water potential. This dependence on water potential is best described phenomenologically by a Weibull function,

$$\kappa(h,\psi) = \kappa_{\max}(h)P(h,\psi) = \kappa_{\max}(h)\left(\frac{1}{2}\right)^{\left(\frac{\psi}{\psi_{50}(h)}\right)^{b}},$$

where $\kappa_{\max}(\mathrm{h})$ is the maximum conductivity at distance $h$ (i.e., the conductivity at $\psi = 0$), $P(h,\psi)$ is the so-called vulnerability curve (Eq. 2 in the main text) of a slice of the conducting segment at distance $h$, $\psi_{50}(\mathrm{h})$ is the water potential at which 50% conductivity is lost at distance $h$ (i.e., $\kappa\big(h,\psi_{50}(\mathrm{h})\big) = \kappa_{\max}(\mathrm{h})/2$), and $b$ is the vulnerability curve's shape parameter, such that a higher value of $b$ leads to a steeper decline in conductivity around $\psi = \psi_{50}(\mathrm{h})$. Note that both $\kappa_{\max}(\mathrm{h})$ and $\psi_{50}(\mathrm{h})$ can in general depend on the distance $h$ from the plant base due to geometric changes of pipes along the hydraulic pathway as described above.

##### 1.1.4 Equation governing water flow

The drop of water potential along a hydraulic pathway's cross-section over a small vertical distance increment $\mathrm{d}h$ occurs for two reasons, due to gravity and due to water flow. The drop in water potential due to gravity is $\rho g_{\parallel}\mathrm{d}h$ (first term on the right-hand side in the equation below), where $\rho$ is the density of water and $g_{\parallel}$ is the component of the gravitational acceleration in the direction of the water flow. The drop due to a volumetric flow rate of liquid water per unit area, $Q$, through the pathway follows from the already mentioned Darcy's law and the relation between conductivity and conductance per unit area (second term on the right-hand side in the equation below). Thus, the total drop in water potential is given by (Fig. M1),

$$\psi(h+\mathrm{d}h) - \psi(h) = -\rho g_{\parallel}\mathrm{d}h - \frac{Q\eta\mathrm{d}h}{\kappa\big(h,\psi(h)\big)},$$

where $\psi(h)$ is the water potential at a distance $h$ from the plant base, $\mathrm{d}h$ is the considered small distance increment, $\eta$ is the dynamic viscosity of water, and $\kappa\big(h,\psi(h)\big)$ is the conductivity at a distance $h$ with water potential $\psi(h)$.

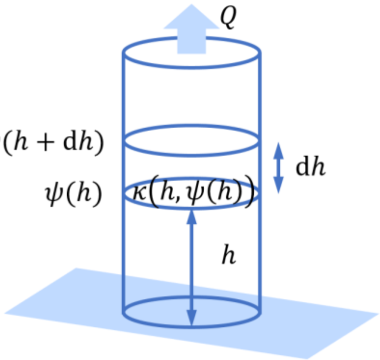

**Fig. M1.** Schematic illustration of the flow of water through a conducting segment.

This yields a differential equation describing the water flow through any conducting segment whose conductivity may vary with distance and water potential,

$$\frac{\mathrm{d}\psi}{\mathrm{d}h} = -\rho g_{\parallel} - \frac{Q\eta}{\kappa_{\max}(h) P\big(h, \psi(h)\big)}, \qquad \text{S1}$$

##### 1.1.5 Solution of the equation governing water flow

If we ignore the effects of gravity (assuming $\rho g_{\parallel} \ll Q\eta/\kappa$) and xylem tapering (assuming $\kappa_{\max}(h)$ to be independent of $h$ and $P\big(h, \psi(h)\big)$ to depend on $h$ only through $\psi(h)$), we can solve Eq. S1 to express $Q$ as

$$Q = -\frac{\kappa_{\max}}{L\eta} \int_{\psi_{\mathrm{s}}}^{\psi_{\mathrm{l}}} P(\psi)\mathrm{d}\psi\,, \qquad \text{S2}$$

where $\psi_{\mathrm{s}}$ and $\psi_{\mathrm{l}}$ are the water potentials in the soil and in the leaves (near the stomata), respectively, and $L$ is the length of the hydraulic pathway between the soil and the stomata. The vulnerability curve $P(\psi)$ here describes the combined effect of the loss of conductivity of the stem and leaf xylem, as well as of the outside-xylem segment.

The parameter $K_{\mathrm{L}}$ estimated in our model fits is maximum leaf-specific whole-plant conductance, i.e., the maximum conductance of the entire plant including roots, stem, and leaves, expressed per unit leaf area. It equals the ratio $\kappa_{\max}/L$, and represents the combined effect of xylem and leaf permeabilities, plant height, and leaf thickness.

##### 1.1.6 Transpiration flow

Water exits from the leaves due to transpiration. The volumetric transpiration flow rate $E$ of water vapour per unit leaf area is given by Fick's law of diffusion,

$$E = 1.6 g_{\mathrm{s}}(e_{\mathrm{l}} - e_{\mathrm{a}}),$$

where $g_{\mathrm{s}}$ is the stomatal conductance to $CO_2$, and the factor 1.6 is the ratio of the diffusivities of water vapour and $CO_2$, which therefore converts $g_{\mathrm{s}}$ to the stomatal conductance to water vapour. Furthermore, $e_{\mathrm{a}}$ is the relative partial pressure of water vapour in the atmosphere, and $e_{\mathrm{l}}$ is the relative partial pressure of water vapour inside the leaf, both of which are expressed as fractions of the atmospheric pressure.

The air inside the leaf is typically saturated with water vapour, such that $e_\mathrm{l} \approx e_\mathrm{s}$, where $e_\mathrm{s}$ is the saturation vapour pressure as a fraction of the atmospheric pressure, and the difference $e_\mathrm{s} - e_\mathrm{a}$ is the atmospheric vapour pressure deficit $D$. Thus,

$$E \approx 1.6 g_s D. \qquad \text{S3}$$

Notice that, even though $g_\mathrm{s}$ is typically referred to as a "conductance", its definition following from Fick's law of diffusion is that of a diffusion coefficient, rather than that of a hydraulic conductance used for characterizing the flow of liquid water.

##### 1.1.7 Water-balance principle

Since there is no water storage in the leaves, water balance applies. This means that the hydraulic flow rate $Q$ at which water enters the leaves according to Eq. S2 equals the transpiration flow rate $E$ at which water vapour diffuses out of the leaves into the atmosphere according to Eq. S3,

$$Q = E.$$

This allows us to express $g_\mathrm{s}$ in terms of $\Delta\psi = \psi_\mathrm{s} - \psi_\mathrm{l}$,

$$g_\mathrm{s} = -\frac{K_\mathrm{L}}{1.6 D \eta} \int_{\psi_\mathrm{s}}^{\psi_\mathrm{s} - \Delta\psi} P(\psi)\,\mathrm{d}\psi\,, \qquad \text{S4}$$

which shows that, as a consequence of water balance, $g_\mathrm{s}$ is independent of $\chi$.

| Symbol | Meaning | Unit |
|---|---|---|
| $k$ | Conductance | $m^3$ |
| $K$ | Conductance per unit area | m |
| $\kappa$ | Conductivity, i.e., conductance per unit area at unit length | $m^2$ |
| $\psi$ | Water potential | Pa |
| $\psi_{50}$ | Water potential at which 50% conductivity is lost | Pa |
| $\psi_s$ | Soil water potential | Pa |
| $\psi_\mathrm{l}$ | Leaf water potential | Pa |
| $\eta$ | Viscosity of water | Pa s |
| $\rho$ | Density of water | $kg\ m^{-3}$ |
| $P(\psi)$ | Vulnerability curve, i.e., fraction of maximum conductivity remaining at water potential $\psi$ | – |
| $q$ | Volumetric flow rate of liquid water | $m^3\ s^{-1}$ |
| $Q$ | Volumetric flow rate of liquid water per unit area | $m\ s^{-1}$ |
| $E$ | Volumetric transpiration flow rate of water vapour per unit area | $m\ s^{-1}$ |
| $g_s$ | Stomatal conductance, i.e., amount of $CO_2$ or water vapour entering or exiting leaves per unit leaf area per unit concentration gradient | $mol\ m^{-2}\ s^{-1}$ |
| $e_\mathrm{a}$ | Partial pressure of water vapour in the atmosphere divided by atmospheric pressure | – |

| | | |
|---|---|---|
| $e_l$ | Partial pressure of water vapour inside the leaf divided by atmospheric pressure | – |
| $e_s$ | Saturation vapour pressure divided by atmospheric pressure | – |
| $D$ | Vapour pressure deficit divided by atmospheric pressure | – |

**Table M1. Variables and notations in the water-transport module.**

#### 1.2 Photosynthesis module and photosynthetic-coordination hypothesis

In this section, we describe the standard biochemical model of photosynthesis, also called the Farquhar-von Caemmerer-Berry biochemical model, and specify and apply the photosynthetic-coordination hypothesis.

##### 1.2.1 Standard biochemical model of photosynthesis

Photosynthesis is the process by which plant leaves use light energy to convert $CO_2$ from the atmosphere and water from the soil into sugars. It depends on two sequential reactions: (1) the light-dependent electron-transport reaction, through which photosystems in the leaf use light energy to strip electrons from molecules of water and use them to produce NADPH and ATP molecules, which act as short-term energy reserves, and (2) the light-independent carboxylation reaction, through which NADPH and ATP molecules are used to power the Calvin cycle, in which $CO_2$ is absorbed with the help of an enzyme called RuBisCO and fixed into sugars. Thus, photosynthesis is jointly limited by light availability (which is further subject to the efficiency of the photosystems) and the carboxylation capacity of leaves (i.e., the chemical activity of RuBisCO).

The carboxylation-limited rate $A_\mathrm{c}$ of photosynthesis is given by the Farquhar-von Caemmerer-Berry biochemical model (Farquhar et al., 1980) ,

$$A_c = V_\mathrm{cmax}\frac{c_\mathrm{i} - \Gamma^*}{c_\mathrm{i} + K_\mathrm{M}} - R_d, \qquad \text{S5}$$

where $V_\mathrm{cmax}$ is the maximum carboxylation capacity of leaves, $c_\mathrm{i}$ is the leaf-internal $CO_2$ concentration, $\Gamma^*$ is the photosynthetic light-compensation point, $K_\mathrm{M}$ is the Michaelis-Menten coefficient for C3 photosynthesis, and $R_\mathrm{d}$ is the dark-respiration rate, which is assumed to be proportional to $V_\mathrm{cmax}$,

$$R_\mathrm{d} = b_\mathrm{r} V_\mathrm{cmax},$$

where $b_\mathrm{r}$ is the dark-respiration rate per unit carboxylation capacity. The quantities $c_\mathrm{i}$ , $\Gamma^*$, and $K_\mathrm{M}$ are all expressed as partial pressures divided by the atmospheric pressure and are thus unitless.

The light-limited rate $A_\mathrm{j}$ of photosynthesis is also given by the Farquhar-von Caemmerer-Berry biochemical model,

$$A_\mathrm{j} = \frac{J}{4}\left(\frac{c_\mathrm{i} - \Gamma^*}{c_\mathrm{i} + 2\Gamma^*}\right) - R_\mathrm{d}, \qquad \text{S6}$$

where $J$ is the effective electron-transport capacity of the leaves, jointly determined by the light availability $I_\mathrm{abs}$ and the leaves' intrinsic maximum electron-transport capacity $J_\mathrm{max}$. The factor '4' accounts for the fact that for every four photons absorbed by the leaf's photosystems, one electron is released. The response of $J$ to light availability is saturating, phenomenologically expressed by a rectangular hyperbola,

$$J = \frac{4\phi_0 I_{\mathrm{abs}}}{\sqrt{\left(\frac{4\phi_0 I_{\mathrm{abs}}}{J_{\mathrm{max}}}\right)^2 + 1}}, \qquad \text{S7}$$

where $J_{\mathrm{max}}$ is the maximum electron-transport capacity of leaves (approached when $I_{\mathrm{abs}} \to \infty$), $\phi_0$ is the intrinsic quantum-yield efficiency, and $I_{\mathrm{abs}}$ is the incident photosynthetic photon-flux density, measuring light availability in terms of the absorbed photosynthetically active solar radiation.

Inverting the above relation, we can express $J_{\mathrm{max}}$ in terms of $J$,

$$J_{\mathrm{max}} = \frac{4\phi_0 I_{\mathrm{abs}}}{\sqrt{\left(\frac{4\phi_0 I_{\mathrm{abs}}}{J}\right)^2 - 1}}. \qquad \text{S8}$$

Finally, since the carboxylation reaction follows electron-transport reaction, the rate $A_{\mathrm{p}}$ of photosynthesis is limited by the slower of the two reactions,

$$A_{\mathrm{p}} = \min\left(\mathrm{A_c}, \mathrm{A_j}\right).$$

##### 1.2.2 Rate of $CO_2$ diffusion into the leaves

The rate $A_{\mathrm{d}}$ at which $CO_2$ diffuses from the atmosphere into the leaves is again given by Fick's law,

$$A_{\mathrm{d}} = g_{\mathrm{s}}(c_{\mathrm{a}} - c_{\mathrm{i}}) = g_{\mathrm{s}} c_{\mathrm{a}}(1 - \chi), \qquad \text{S9}$$

where $\chi$ is the ratio $c_{\mathrm{i}}/c_{\mathrm{a}}$ of the leaf-internal and ambient $CO_2$ partial pressures.

##### 1.2.3 $CO_2$ balance

Under equilibrium conditions, there is no storage or build-up of $CO_2$ in the leaves. Therefore, the rate of photosynthesis, i.e., the rate at which $CO_2$ is fixed by the leaves, equals the rate at which $CO_2$ diffuses into the leaves,

$$A_{\mathrm{d}} = A_{\mathrm{p}}.$$

##### 1.2.4 Photosynthetic-coordination hypothesis

The photosynthetic-coordination hypothesis (see main text) states that, under typical daytime conditions, the photosynthetic capacities are modulated by leaves such that the carboxylation-limited and light-limited assimilation rates are equal,

$$A_{\mathrm{c}} = A_{\mathrm{j}}.$$

Therefore,

$$V_{\mathrm{cmax}} \frac{c_{\mathrm{i}} - \Gamma^*}{c_{\mathrm{i}} + K_{\mathrm{M}}} - R_{\mathrm{d}} = \frac{J}{4}\left(\frac{c_{\mathrm{i}} - \Gamma^*}{c_{\mathrm{i}} + 2\Gamma^*}\right) - R_{\mathrm{d}},$$

which gives

$$V_{\mathrm{cmax}} = \frac{J}{4}\left(\frac{c_{\mathrm{i}} + K_{\mathrm{M}}}{c_{\mathrm{i}} + 2\Gamma^*}\right). \quad \text{S10}$$

Hence, $A_{\mathrm{j}}$ can be rewritten as

$$A_{\mathrm{j}} = \frac{J}{4} \cdot \frac{c_{\mathrm{i}}(1 - b_{\mathrm{r}}) - (\Gamma^* + b_{\mathrm{r}} K_{\mathrm{M}})}{c_{\mathrm{i}} + 2\Gamma^*}. \quad \text{S11}$$

Equating Eq. S9 and Eq. S11, $J$ can be expressed in terms of $\chi$ as

$$J = 4 g_{\mathrm{s}} c_{\mathrm{a}} \frac{(1 - \chi)(\chi c_{\mathrm{a}} + 2\Gamma^*)}{\chi c_{\mathrm{a}}(1 - b_{\mathrm{r}}) - (\Gamma^* + b_{\mathrm{r}} K_{\mathrm{M}})}. \quad \text{S12}$$

By substituting Eq. S12 into Eq. S8, we obtain an expression for $J_{\mathrm{max}}$ in terms of $g_{\mathrm{s}}$ and $\chi$.

| Symbol | Meaning | Unit |
|---|---|---|
| $A_{\mathrm{c}}$ | Carboxylation-limited photosynthesis rate | mol $m^{-2}$ $s^{-1}$ |
| $A_{\mathrm{j}}$ | Electron-transport-limited photosynthesis rate | mol $m^{-2}$ $s^{-1}$ |
| $A_{\mathrm{p}}$ | Rate of photosynthesis, i.e., rate of $CO_2$ fixation in the leaves | mol $m^{-2}$ $s^{-1}$ |
| $A_{\mathrm{d}}$ | Rate at which $CO_2$ diffuses from the atmosphere into the leaves | mol $m^{-2}$ $s^{-1}$ |
| $c_{\mathrm{i}}$ | Leaf-internal $CO_2$ partial pressure, divided by atmospheric pressure | – |
| $c_{\mathrm{a}}$ | Ambient $CO_2$ partial pressure, divided by atmospheric pressure | – |
| $\chi$ | Ratio of $c_{\mathrm{i}}$ and $c_{\mathrm{a}}$ | – |
| $\Gamma^*$ | Photosynthetic light-compensation point | – |
| $K_{\mathrm{M}}$ | Michaelis-Menten coefficient for C3 photosynthesis | – |
| $V_{\mathrm{cmax}}$ | Maximum carboxylation capacity of leaves | mol $m^{-2}$ $s^{-1}$ |
| $J$ | Effective electron-transport capacity of leaves | mol $m^{-2}$ $s^{-1}$ |
| $J_{\mathrm{max}}$ | Maximum electron-transport capacity of leaves, under light saturation | mol $m^{-2}$ $s^{-1}$ |
| $R_{\mathrm{d}}$ | Dark-respiration rate | mol $m^{-2}$ $s^{-1}$ |
| $\phi_0$ | Intrinsic quantum-yield efficiency | – |
| $I_{\mathrm{abs}}$ | Incident photosynthetic photon-flux density | mol $m^{-2}$ $s^{-1}$ |
| $b_{\mathrm{r}}$ | Dark-respiration rate per unit carboxylation capacity | – |

**Table M2. Variables and notations in the photosynthesis module.**

#### 1.3 Optimization module and profit-maximization hypothesis

In this section, we specify and apply the profit-maximization hypothesis and describe our semi-analytical solution to the resulting optimality problem, as well as a useful special case possessing a simple analytical solution.

##### 1.3.1 Profit-maximization hypothesis

We assume that plants maximize their profit $F$ defined as the benefit from photosynthetic assimilation minus the costs of maintaining the photosynthetic capacities and the hydraulic pathway,

$$F = A_{\mathrm{j}} - \alpha J_{\max} - \gamma \Delta\psi^2,$$

where each term depends on at least one of the two independent variables $\chi$ and $\Delta\psi$,

$$F(\chi, \Delta\psi) = g_{\mathrm{s}}(\Delta\psi) c_{\mathrm{a}}(1-\chi) - \alpha J_{\max}(\chi, \Delta\psi) - \gamma \Delta\psi^2.$$

##### 1.3.2 Semi-analytical solution to the profit-maximization model

To solve the optimality condition, we set the gradient of the profit function to zero,

$$\begin{aligned} \frac{\partial F}{\partial \chi} &= -g_{\mathrm{s}} c_{\mathrm{a}} - \alpha \frac{\partial J_{\max}}{\partial J} \frac{\partial J}{\partial \chi} - 0 = 0, \\ \frac{\partial F}{\partial \Delta\psi} &= \frac{\partial g_{\mathrm{s}}}{\partial \Delta\psi} c_{\mathrm{a}}(1-\chi) - \alpha \frac{\partial J_{\max}}{\partial J} \frac{\partial J}{\partial \Delta\psi} - 2\gamma \Delta\psi = 0. \end{aligned} \qquad \text{S13}$$

The four derivatives required in the above equations are as follows,

$$\frac{\partial J_{\max}}{\partial J}=\frac{(4\phi_0 I_{\rm abs})^3}{((4\phi_0 I_{\rm abs})^2-J^2)^{3/2}},$$
$$\frac{\partial J}{\partial \chi}=4g_{\rm s}c_{\rm a}\left(b_{\rm r}\frac{2\frac{\Gamma^*}{c_{\rm a}}\left(\frac{K_{\rm M}}{c_{\rm a}}+1\right)+\frac{K_{\rm M}}{c_{\rm a}}(2\chi-1)+\chi^2}{\left(b_{\rm r}\left(\frac{K_{\rm M}}{c_{\rm a}}+\chi\right)+\frac{\Gamma^*}{c_{\rm a}}-\chi\right)^2}-\frac{(\chi-\frac{\Gamma^*}{c_{\rm a}})^2+3\frac{\Gamma^*}{c_{\rm a}}(1-\frac{\Gamma^*}{c_{\rm a}})}{\left(b_{\rm r}\left(\frac{K_{\rm M}}{c_{\rm a}}+\chi\right)+\frac{\Gamma^*}{c_{\rm a}}-\chi\right)^2}\right), \quad \text{S14}$$
$$\frac{\partial J}{\partial \Delta\psi}=4\frac{\partial g_{\rm s}}{\partial \Delta\psi}c_{\rm a}\frac{(1-\chi)(\chi c_{\rm a}+2\Gamma^*)}{\chi c_{\rm a}(1-b_{\rm r})-(\Gamma^*+b_{\rm r}K_{\rm M})},$$
$$\frac{\partial g_{\rm s}}{\partial \Delta\psi}=\frac{\partial}{\partial \Delta\psi}\left(-\frac{K_{\rm L}}{1.6D\eta}\int_{\psi_{\rm s}}^{\psi_{\rm s}-\Delta\psi}P\,(\psi)\mathrm{d}\psi\right)=\frac{K_{\rm L}}{1.6D\eta}P(\psi_{\rm s}-\Delta\psi).$$

By substituting the derivatives in Eq. S14 into Eq. S13, we obtain two equations involving the unknowns $\chi$ and $\Delta\psi$, which could be solved using a two-dimensional root-finding algorithm.

However, Eq. S13 can be simplified analytically: to this end, we first rearrange both equations in Eq. S13 to move $\partial J_{\max}/\partial J$ to their left-hand sides and then eliminate this term by equating the resultant right-hand sides,

$$-\frac{g_{\rm s}c_{\rm a}}{\alpha\frac{\partial J}{\partial \chi}}=\frac{g_{\rm s}'(\Delta\psi)c_{\rm a}(1-\chi)-2\gamma\Delta\psi}{\alpha\frac{\partial J}{\partial \Delta\psi}}. \quad \text{S15}$$

This equation yields $\chi$ in terms of $\Delta\psi$,

$$\chi^*(\Delta\psi)$$
$$=\Big(c_{\rm a}^2\big((3-2b_{\rm r})\Gamma^*+b_{\rm r}K_{\rm M}\big)g_{\rm s}'(\Delta\psi)-2c_{\rm a}\Delta\psi\gamma(b_{\rm r}K_{\rm M}+\Gamma^*)$$
$$-\sqrt{2c_{\rm a}^2\Delta\psi\gamma\big((2b_{\rm r}-3)\Gamma^*-b_{\rm r}K_{\rm M}\big)\big((b_{\rm r}-1)c_a+b_{\rm r}K_{\rm M}+\Gamma^*\big)\big((c_{\rm a}+2\Gamma^*)g_{\rm s}'(\Delta\psi)-2\Delta\psi\gamma\big)}\Big)$$
$$/\Big(c_{\rm a}^2\Big(\big((3-2b_{\rm r})\Gamma^*+b_{\rm r}K_{\rm M}\big)g_{\rm s}'(\Delta\psi)+2(b_{\rm r}-1)\Delta\psi\gamma\Big)\Big).$$

Substituting this solution for $\chi$ into Eq. S13 results in all terms to depend on $\Delta\psi$ alone. The optimal $\Delta\psi^*$ can then be found using a one-dimensional root-finding algorithm, instead of solving Eqs. S13 and S14 using a more complex two-dimensional root-finding algorithm.

##### 1.3.3 Fully analytical solution for a special case

In the special case of strong electron-transport limitation or, equivalently, high light availability, the optimal $\chi$ can be obtained analytically, and turns out to be independent of $\Delta\psi$: $J_{\max}\ll 4\phi_0 I_{\rm abs}$ implies $J\approx J_{\max}$ and thus

$$\frac{\partial J_{\max}}{\partial J} = 1,$$

from which we obtain

$$\frac{\partial F}{\partial \chi} = -g_{\mathrm{s}} c_{\mathrm{a}} - \alpha \frac{\partial J}{\partial \chi} = 0.$$

This is a quadratic equation in $\chi$, giving

$$\chi^* = \frac{(1 - 4\alpha - b_{\mathrm{r}})\left(b_{\mathrm{r}} K_{\mathrm{M}} + \frac{\Gamma^*}{c_{\mathrm{a}}}\right) + \sqrt{4\alpha(1 - 4\alpha - b_{\mathrm{r}})\left(3\frac{\Gamma^*}{c_{\mathrm{a}}} - b_{\mathrm{r}}\left(2\frac{\Gamma^*}{c_{\mathrm{a}}} + K_{\mathrm{M}}\right)\right)\left(1 - \frac{\Gamma^*}{c_{\mathrm{a}}} - b_{\mathrm{r}}(1 + K_{\mathrm{M}})\right)}}{(1 - b_{\mathrm{r}})(1 - 4\alpha - b_{\mathrm{r}})}. \qquad S16$$

In the absence of dark respiration ($b_{\mathrm{r}} = 0$), this simplifies to

$$\chi^* = \frac{\frac{\Gamma^*}{c_{\mathrm{a}}}(1 - 4\alpha) + \sqrt{12\alpha(1 - 4\alpha)\frac{\Gamma^*}{c_{\mathrm{a}}}\left(1 - \frac{\Gamma^*}{c_{\mathrm{a}}}\right)}}{1 - 4\alpha}.$$

| Symbol | Meaning | Unit |
|---|---|---|
| $F$ | Plant profit from photosynthesis | mol m$^{-2}$ s$^{-1}$ |
| $\alpha$ | Unit cost of photosynthetic capacity | – |
| $\gamma$ | Unit cost of hydraulic capacity | mol m$^{-2}$ s$^{-1}$ Pa$^{-2}$ |

**Table M3. Variables and notations in the optimization module.**

### 2 Model extension

In the present study, we use an unsegmented water-transport model, in which the entire hydraulic pathway is represented by a single effective segment, as shown in Section 1.1. To facilitate future research, we also derive and present an extended two-segment version of the water-transport model that can explicitly account for the hydraulic properties of the stem and the leaves, as shown in Fig. 1.

#### 2.1 Extended water-transport module and water-balance principle

To derive the total drop in water potential from the soil to the air, we must find and add the drops in water potential along the xylem and outside-xylem segments. These two segments approximately correspond to locations in the stems and the leaves of plants, respectively.

##### 2.1.1 Drop in water potential across the xylem segment

The Hagen-Poiseuille equation describes the pressure drop $\Delta\psi$ along a cylindrical pipe of length $L$ and radius $R$ due to water flowing in it at the rate $q$,

$$\Delta\psi = \frac{8\eta L q}{\pi R^4},$$

where $\eta$ is the dynamic viscosity of water.

Water is conducted through the stem and leaf veins by interconnected xylem vessels. Using Darcy's law as stated in Section 1.1.2, the conductance $k_{\mathrm{v}}$ of a single cylindrical vessel of length $l_{\mathrm{v}}$ and radius $r_{\mathrm{v}}$ thus equals

$$k_{\mathrm{v}} = \frac{\eta q}{\Delta\psi} = \frac{\pi r_{\mathrm{v}}^4}{8 l_{\mathrm{v}}}.$$

If the sapwood has a vessel density $\rho_{\mathrm{v}}$, measuring the number of parallel vessels per unit sapwood area, vessel conductances will add up such that the conductivity $\kappa_{\mathrm{s}}$ of the sapwood xylem is given by

$$\kappa_{\mathrm{s}} = k_{\mathrm{v}} l_{\mathrm{v}} \rho_{\mathrm{v}} = \frac{\pi \rho_{\mathrm{v}} r_{\mathrm{v}}^4}{8}.$$

For example, in *Austromyrtus bidwillii*, $\rho_{\mathrm{v}} = 217\ \mathrm{mm}^{-2}$ and $r_{\mathrm{v}} = 15.25\ \mu\mathrm{m}$ (Choat et al., 2005), giving $\kappa_{\mathrm{s}} = 4.6 \times 10^{-12}\ \mathrm{m}^2$. The conductance of the xylem segment per unit sapwood area is given by $K_{\mathrm{s}} = k_{\mathrm{v}} \rho_{\mathrm{v}} = \kappa_{\mathrm{s}} / l_{\mathrm{v}}$.

The ratio of sapwood area to leaf area is known as the Huber value $\nu_{\mathrm{H}}$. When applying Eq. S1 to plants, it is convenient to define $Q$ as the rate of volumetric water flow per unit leaf area rather than per unit sapwood area. Therefore, we multiply the sapwood conductivity $\kappa_{\mathrm{s}}$ by $\nu_{\mathrm{H}}$ to obtain the sapwood conductivity per unit leaf area.

Again ignoring the effects of gravity and xylem tapering as in Section 1.1.5, we can solve Eq. S1 for the xylem segment to express water flux $Q_{\mathrm{x}}$ in the xylem-segment as

$$Q_{\mathrm{x}} = -\frac{\kappa_{\mathrm{s}} \nu_{\mathrm{H}}}{L_{\mathrm{x}} \eta} \int_{\psi_{\mathrm{s}}}^{\psi_{\mathrm{p}}} P_{\mathrm{x}}(\psi) \mathrm{d}\psi, \qquad \text{S17}$$

where $\psi_{\mathrm{s}}$ and $\psi_{\mathrm{p}}$ are the water potentials in the soil and at the end of the leaf veins, respectively, $L_{\mathrm{x}}$ is the length of the xylem pathway to which plant height is the key contributor, and $P_{\mathrm{x}}$ is the vulnerability curve of the xylem, in which the loss of conductivity occurs largely through cavitation.

##### 2.1.2 Drop in water potential across the outside-xylem segment

Once the water exits the xylem at the end of the leaf veins, it passes through bundle sheath cells and several layers of spongy mesophyll cells before reaching the stomata, from where it vaporizes from the cell walls and out into the atmosphere. This outside-xylem segment often offers much larger resistance to water flow than the xylem segment and lose conductivity much faster than the xylem segment as water potential decreases, making the leaves the hydraulic bottleneck in a plant's flow of water.

We can represent the total flow $Q_{\mathrm{ox}}$ through all different outside-xylem hydraulic structures (including the cell walls and the cell mesophyll) in the same way as above,

$$Q_{\mathrm{ox}} = -\frac{\kappa_{\mathrm{l}}}{L_{\mathrm{ox}} \eta} \int_{\psi_{\mathrm{p}}}^{\psi_{\mathrm{l}}} P_{\mathrm{ox}}(\psi) \mathrm{d}\psi, \qquad \text{S18}$$

where $\kappa_\mathrm{l}$ is the conductivity of leaves per unit leaf area, $\psi_\mathrm{l}$ is the water potential in the leaves (near the stomata), and $L_\mathrm{ox}$ is the length of the hydraulic pathway outside the xylem, which depends on leaf thickness and leaf venation density.

##### 2.1.3 Flow continuity between xylem and outside-xylem segments

Since there is no loss or storage of water at the interface of the xylem and outside-xylem segments of the hydraulic pathway, the rate $Q_\mathrm{x}$ at which water flows in the xylem segment equals the rate $Q_\mathrm{ox}$ at which it flows in the outside-xylem segment, $Q = Q_\mathrm{x} = Q_\mathrm{ox}$. Substituting these rates according to Eqs. S17 and S18, we obtain

$$Q = -\frac{\kappa_\mathrm{s} \nu_\mathrm{H}}{L_\mathrm{x} \eta} \int_{\psi_\mathrm{s}}^{\psi_\mathrm{s} - \Delta\psi_\mathrm{x}} P_\mathrm{x}(\psi) \mathrm{d}\psi = -\frac{\kappa_\mathrm{l}}{L_\mathrm{ox} \eta} \int_{\psi_\mathrm{s} - \Delta\psi_\mathrm{x}}^{\psi_\mathrm{s} - \Delta\psi} P_\mathrm{ox}(\psi) \mathrm{d}\psi, \qquad \text{S19}$$

where the integration limits follow from that fact that the drops in water potential across the xylem and outside-xylem segments are $\Delta\psi_\mathrm{x} = \psi_\mathrm{s} - \psi_\mathrm{p}$ and $\Delta\psi_\mathrm{ox} = \psi_\mathrm{p} - \psi_\mathrm{l}$, respectively, and the total soil-to-leaf difference in water potential is $\Delta\psi = \Delta\psi_\mathrm{x} + \Delta\psi_\mathrm{ox} = \psi_\mathrm{s} - \psi_\mathrm{l}$. For a given $\Delta\psi$, Eq. S19 can be solved for $\Delta\psi_\mathrm{x}$, and either of the two equal expressions in Eq. S19 can then be used to obtain $Q$.

##### 2.1.4 Water-balance principle

As in Section 1.1.7, since there is no water storage in the leaves, water balance applies. This means that the hydraulic flow rate $Q$ at which water enters the leaves according to Eq. S19 equals the transpiration flow rate $E$ at which water vapour diffuses out of the leaves into the atmosphere according to Eq. S3,

$$Q = E = 1.6 g_\mathrm{s} D,$$

from which $g_\mathrm{s}$ can be calculated.

| s | Meaning | Unit |
|---|---|---|
| $k_\mathrm{v}$ | Mean conductance of a single xylem vessel | $m^3$ |
| $r_\mathrm{v}$ | Mean radius of a single xylem vessel | m |
| $l_\mathrm{v}$ | Length of a single xylem vessel | m |
| $\rho_\mathrm{v}$ | Number of xylem vessels per unit sapwood area | $m^{-2}$ |
| $\kappa_\mathrm{s}$ | Conductivity of sapwood, expressed per unit sapwood area | $m^2$ |
| $\kappa_\mathrm{l}$ | Conductivity of leaves, expressed per unit leaf area | $m^2$ |
| $\nu_\mathrm{H}$ | Ratio of sapwood area to leaf area, i.e., Huber value | – |
| $L_\mathrm{x}$ | Path length of the xylem segment, primarily driven by plant height | m |
| $L_\mathrm{ox}$ | Path length of the outside-xylem segment, primarily driven by leaf thickness | m |
| $\psi_\mathrm{p}$ | Water potential at the end of the xylem segment | Pa |
| $Q_\mathrm{x}$ | Volumetric flow rate of liquid water per unit area in the xylem segment | $m\ s^{-1}$ |
| $Q_\mathrm{ox}$ | Volumetric flow rate of liquid water per unit area in the outside-xylem segment | $m\ s^{-1}$ |
| $\Delta\psi_\mathrm{x}$ | Drop in water potential across the xylem segment | Pa |
| $\Delta\psi_\mathrm{ox}$ | Drop in water potential across the outside-xylem segment | Pa |

**Table M4. Variables and notations in the extended water-transport module, beyond those already mentioned in Table M1.**

### 3 Supplementary figures

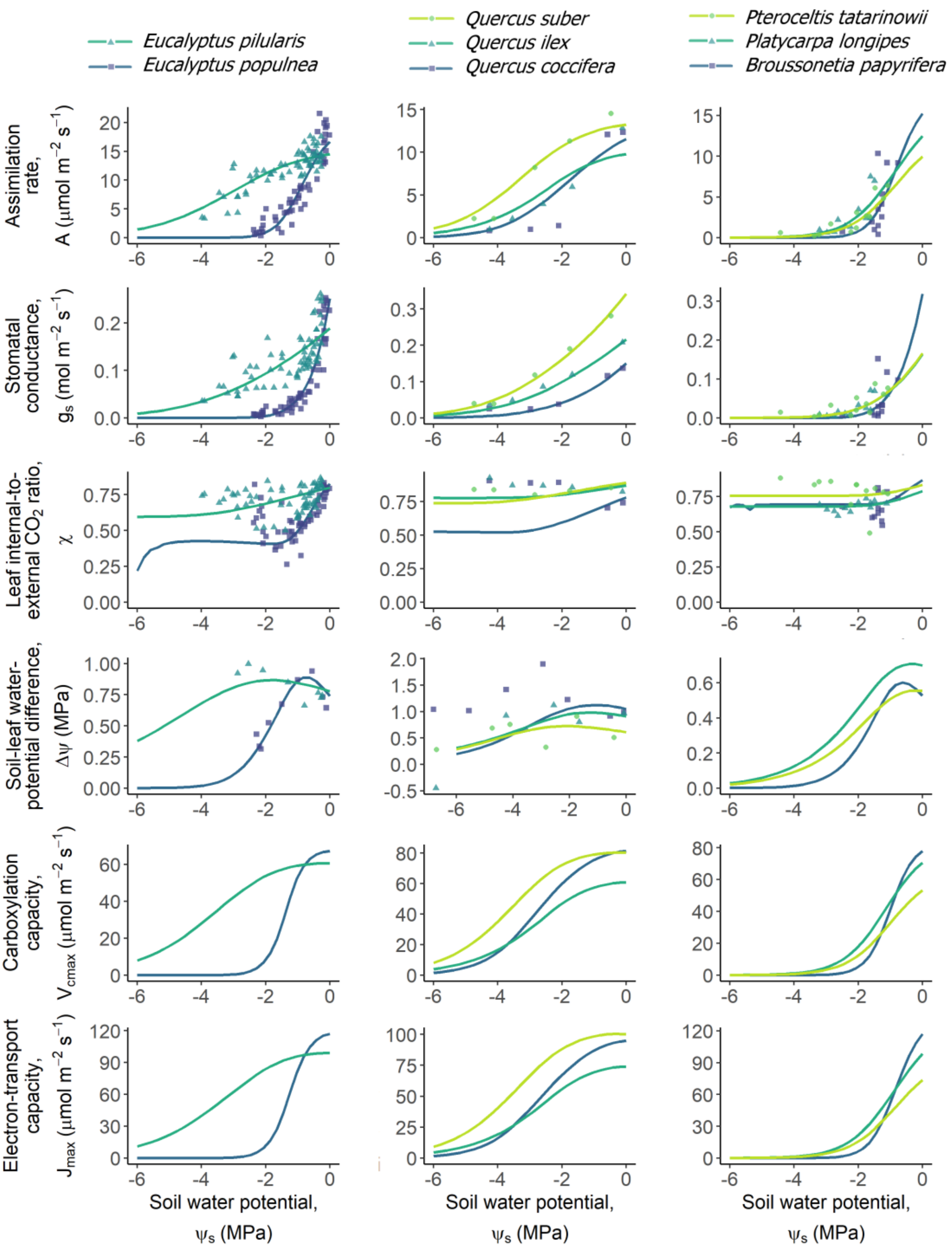

**Fig. S1. Model-predicted and observed responses of the 18 analysed species to soil dry-down.** Our model predictions (lines) are compared to observations (points) for all 18 species covered by our analysis. The first column in this figure reproduces Fig. 2 from the main text showing this comparison for two of the 18 species, and the comparisons for all other 16 species are shown analogously in the further columns. Each column represents a grouping of functionally similar species and has its own colour key: the different species indicated by the colours are shown, separately, at the top of each column. The last two rows do not show data points as direct measurements of photosynthetic capacities are not available; the main text provides details on our indirect validation of these predictions.

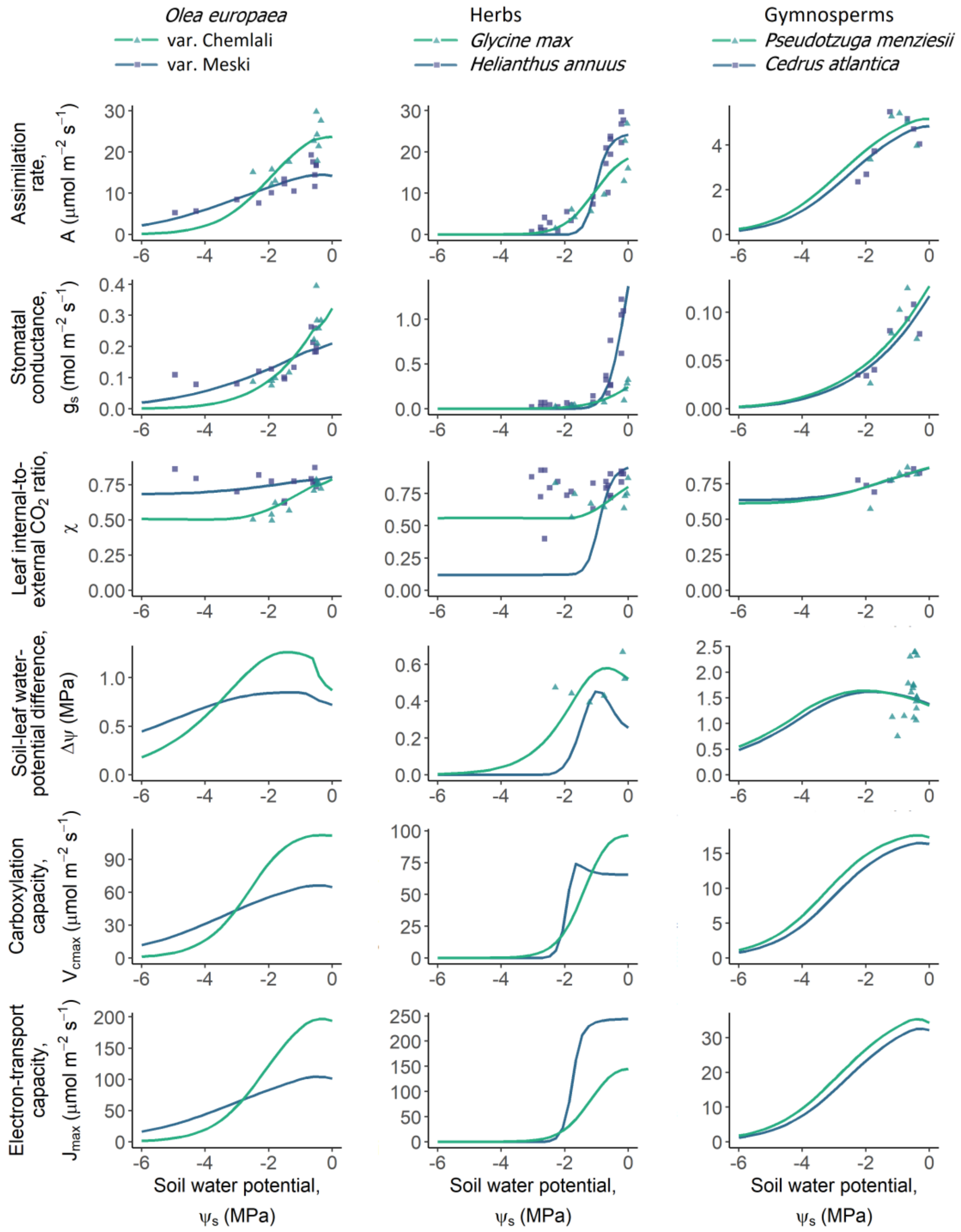

**Fig. S1 (continued).**

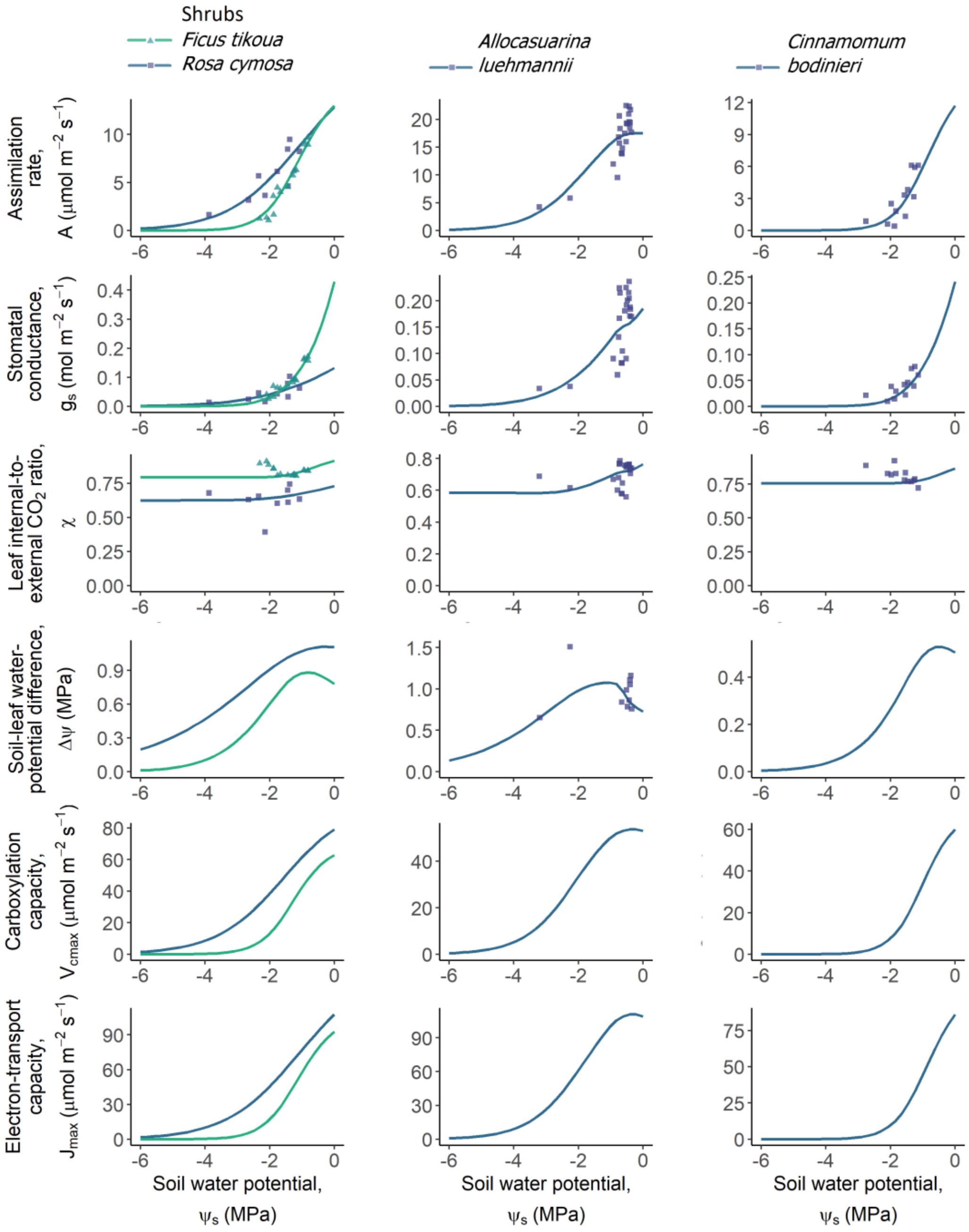

**Fig. S1 (continued).**

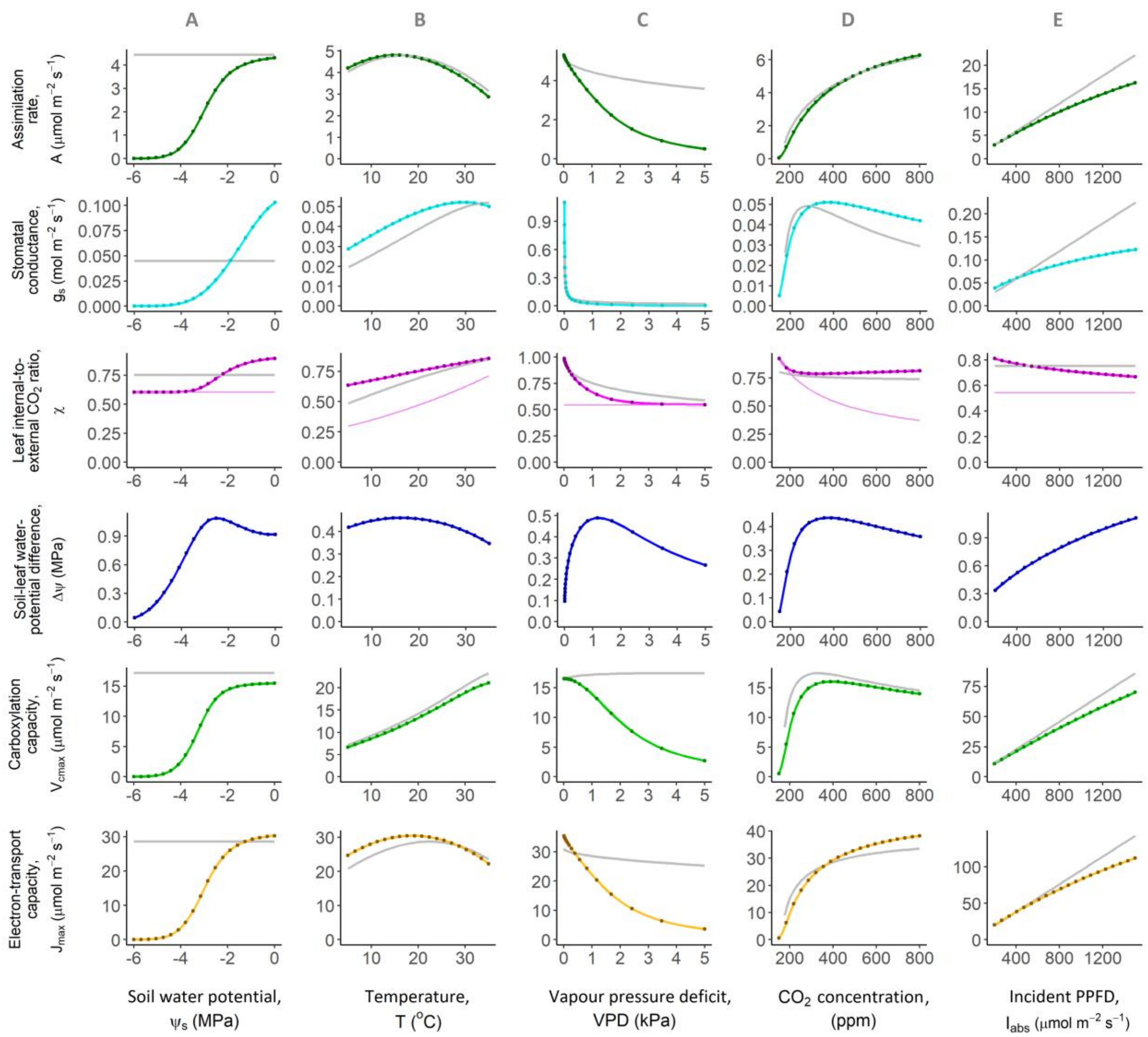

**Fig. S2. Model-predicted responses to atmospheric and soil variables closely resemble those of Wang et al. (2017)**. Photosynthetic responses to atmospheric and soil variables as predicted by our model are shown by coloured lines and compared with those of Wang et al. (2017) shown by grey lines. Here, PPFD is the incident photosynthetic photon-flux density, i.e., light intensity. The model by Wang et al. (2017) lacks an explicit representation of hydraulics (hence no grey lines in the third row) and uses only a single universal cost parameter $\beta$. To facilitate the comparison of our model with theirs, we set $\beta = 146$ in their model and parameterize our model for a plant with the following average parameters: $K_\mathrm{L} = 0.3 \times 10^{-16}$ m$^2$, $\psi_{50} = -2$ MPa, $b = 2$, $\alpha = 0.1$, and $\gamma = 4$. Our model's predictions are shown as obtained from the semi-analytical solution to our model as described in SI Section 1.3.2 (lines) as well as from an independent numerical solution based on numerically maximizing the profit function (points). The accurate match between these semi-analytical and numerical solutions further confirms our numerical results. Thin coloured lines in the fourth row represent the $J_\mathrm{max}$-limited value of $\chi$ determined by Eq. S16.

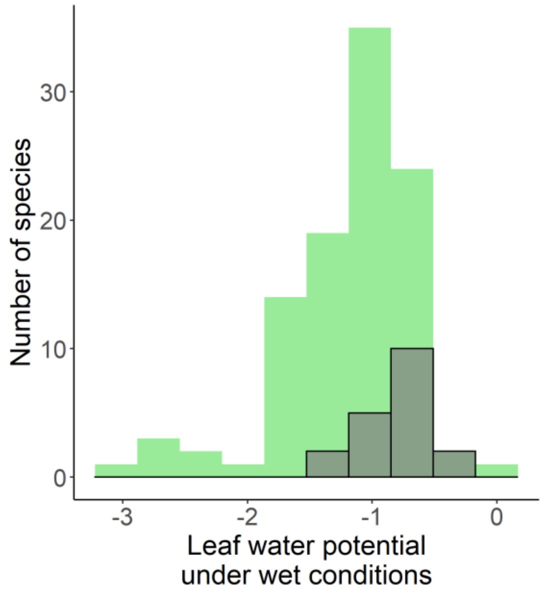

**Fig. S3. Model-predicted and observed frequency distributions of leaf water potentials under wet conditions**. Fig. 5A in the main text showed the predicted and observed distributions of the degree of anisohydricity (slope of the $\psi_l$~$\psi_s$ relationship at $\psi_s = 0$). Similarly, this figure shows the predicted (grey) and observed (green) distributions of the intercept of this relationship, i.e., the leaf water potential under wet conditions. The predicted frequency distribution is broadly consistent with the corresponding distribution observed in global empirical data obtained from Martinez-Vilalta et al. (2014).

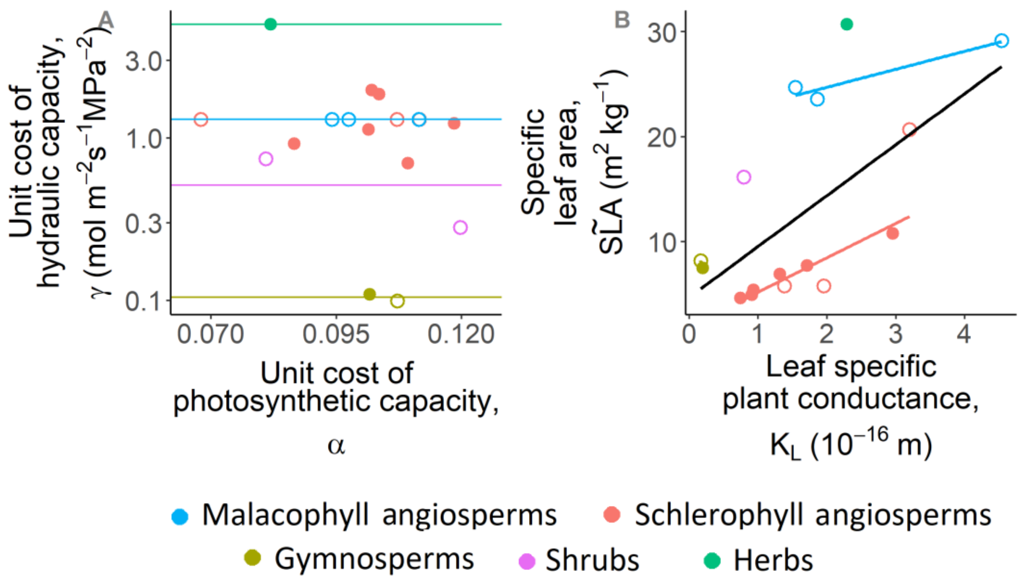

**Fig. S4. Possible simplifications in setting model parameters for global model applications**. (A) The unit costs $\alpha$ of photosynthetic capacity of the 18 analysed species vary within a narrow range, with a value of about 0.1 aptly characterizing most species. Therefore, in global model applications $\alpha$ could be treated as a constant. The unit costs $\gamma$ of hydraulic capacity vary less within than among plant functional types. Therefore, in global model applications $\gamma$ could be treated as constants specific to plant functional types. (B) The plant conductance $K_{\mathrm{L}}$ per unit area is positively correlated with the specific leaf area $S\tilde{L}A$, and this correlation is stronger within than among plant functional types. Therefore, in global model applications $K_{\mathrm{L}}$ could be parameterized using widely available data on specific leaf area in conjunction with linear regressions specific to plant functional types. Closed circles indicate species for which $\gamma$ was estimated using data on $\Delta\psi$, whereas open circles refer to species for which such data was not available and for which we therefore used an average value of $\gamma$ estimated for the respective plant types.

### 4 Cross-validation

| Species | $E_r$ (train) (mean ± st. dev.) | $E_r$ (test) (mean ± st. dev.) | $E_r$ (train) (median) | $E_r$ (test) (median) | Number of data points |
|---|---|---|---|---|---|
| *Allocasuarina luehmannii* | 19.41 ± 2.78 | 24.94 ± 17.26 | 19.69 | 18.27 | 23 |
| *Eucalyptus pilularis* | 30.61 ± 5.05 | 41.18 ± 16.88 | 28.19 | 44.66 | 54 |
| *Eucalyptus populnea* | 14.42 ± 0.49 | 15.53 ± 1.89 | 14.20 | 15.92 | 65 |
| *Glycine max* | 41.58 ± 10.56 | 118.90 ± 154.64 | 41.69 | 48.61 | 9 |
| *Pseudotzuga menziesii* | 21.19 ± 5.27 | 87.62 ± 100.83 | 23.43 | 22.80 | 5 |
| *Quercus coccifera* | 59.32 ± 15.11 | 309.20 ± 347.01 | 53.45 | 134.94 | 6 |
| *Quercus ilex* | 38.10 ± 2.88 | 56.84 ± 16.88 | 39.31 | 49.20 | 5 |
| *Quercus suber* | 17.81 ± 1.86 | 72.00 ± 111.06 | 18.53 | 20.43 | 5 |

**Table S1. Cross-validation results for species with complete data.** We have tested the performance of our model using five-fold cross-validation. For each species for which data was available on the soil-water potential difference $\Delta\psi$, we divided the data on the assimilation rate $A$, the stomatal conductance $g_s$, and the leaf internal-to-external $CO_2$ ratio $\chi$ into five sets. We then performed five training iterations, such that in each iteration we used one of the five sets for testing and the remaining four for training, i.e., estimating parameters. All data on $\Delta\psi$ was used for error calculations. For species with around five data points, each set has only one data point, which makes the algorithm equivalent to leave-one-out cross-validation. The error $E_\mathrm{r}$ is calculated according to Eq. 8. Median errors are comparable across the training and testing datasets, suggesting that our model generalizes well to out-of-sample environmental conditions.

### 5 Sources of empirical trait values

| **Species** | $\tilde{SLA}$ | $\psi_{50x}$ |
|---|---|---|
| *Allocasuarina luehmannii* | TRY | |
| *Helianthus annuus* | TRY | MS |
| *Cedrus atlantica* | TRY | MS |
| *Pseudotzuga menziesii* | BN | MS |
| *Glycine max* | TRY | |
| *Olea europaea* var. Meski | TRY | MA |
| *Olea europaea* var. Chemlali | TRY | MS |
| *Quercus coccifera* | TRY | MA |
| *Quercus suber* | TRY | MS |
| *Broussonetia papyrifera* | L11 | TRY |
| *Eucalyptus pilularis* | AC | |
| *Eucalyptus populnea* | TRY | |
| *Ficus tikoua* | | MS |
| *Cinnamomum bodinieri* | L11 | |
| *Platycarya longipes* | L11 | BA |
| *Pteroceltis tatarinowii* | L11 | BA |
| *Rosa cymosa* | L11 | |
| *Quercus ilex* | TRY | MS |

BN - (Bansal et al., 2015) AC - (Alcorn et al., 2013) MS - (Martin-StPaul et al., 2017) TRY - (Kattge et al., 2011) BA - (Bartlett et al., 2019) MA - (Manzoni et al., 2014)

**Table S2.** Sources of the empirical values of the specific leaf area $S\tilde{L}A$ and the water potential $\psi_{50x}$ at which 50% conductivity is lost in the xylem (Fig. 1).
